# Emergence of cell polarity by reciprocal interactions between Wnt and core PCP components

**DOI:** 10.1101/2025.08.11.669788

**Authors:** Yusuke Mii, Minako Suzuki, Hiroshi Koyama, Kei Nakayama, Ritsuko Takada, Tomoe Kobayashi, Motosuke Tsutsumi, Tomomi Nemoto, Makoto Matsuyama, Toshihiko Fujimori, Shinji Takada

**Affiliations:** National Institute for Basic Biology, National Institutes of Natural Sciences, 5-1 Higashiyama, Myodaiji-cho, Okazaki, Aichi 444-8787, Japan; Exploratory Research Center on Life and Living Systems (ExCELLS), National Institutes of Natural Sciences, 5-1 Higashiyama, Myodaiji-cho, Okazaki, Aichi 444-8787, Japan; The Graduate University for Advanced Studies (SOKENDAI), 5-1 Higashiyama, Myodaiji-cho, Okazaki, Aichi 444-8787, Japan; Institute for Life and Medical Sciences, Kyoto University, 53 Shogoin Kawahara-cho, Sakyo-ku, Kyoto 606-8507, Japan; PRESTO, Japan Science and Technology Agency (JST), Kawaguchi, Saitama, 332-0012, Japan; Kobe Pharmaceutical University, 4-19-1, Motoyamakitamachi, Higashinada-ku, Kobe, 658-8558, Japan; Shigei Medical Research Institutes, 2117 Yamada, Minami-ku, Okayama, 701-0202, Japan; National Institute for Physiological Sciences, National Institutes of Natural Sciences, 5-1 Higashiyama, Myodaiji-cho, Okazaki, Aichi 444-8787, Japan

## Abstract

Planar cell polarity (PCP) is established by asymmetric localization of core PCP components in each cell. Since some Wnt proteins can induce PCP in vertebrates, it has been proposed that Wnt concentration gradients provide spatial cues for polarization. Here, however, we present evidence that Wnt11 can regulate PCP in a gradient-independent manner. In the neural plate of *Xenopus* embryos, endogenous Wnt11 does not form an obvious gradient in the direction of PCP, but is polarized at cell boundaries together with core PCP components. Wnt11 polarization is dependent on core PCP components, while polarization of core PCP components can also be induced by Wnt11, indicating a mutual amplification loop between Wnt11 and core PCP components in PCP formation. Furthermore, Wnt11 and core PCP components compose an intercellular loop to coordinate the direction of polarity. Thus, we propose that local and reciprocal interactions between Wnt11 and core PCP components can generate PCP.

## Introduction

Diffusible signals regulate many biological phenomena, from directional control of migration in single-cell organisms to development and homeostasis of multicellular tissues. In these cases, diffusible signals are thought to act in a top-down manner. For instance, in patterning of developing tissues, it has been thought that these molecules form a concentration gradient around the producing cells, and that positional information provided by the concentration gradient is sensed by receiving cells, thereby orchestrating tissue development (*1–3*). However, it remains unclear whether diffusible signals always exert top-down control over recipient cells, or whether formation of a concentration gradient is actually important in the action of diffusible signals.

Planar cell polarity (PCP) is the collective alignment of individual cell polarities in a tissue plane (*4–10*). A typical example of PCP is observed in *Drosophila* wing, in which the trichome of each cell is aligned in coordinated fashion toward the distal side (*4, 11*). In vertebrates, alignment of stereo-cilia bundles in hair cells in the inner ear (*12*) and orientation of hair follicles in the skin (*13–15*) are examples of PCP. Components essential for establishment of PCP are called core PCP components and include three cell membrane proteins, Fzd (*16*), Vangl (Van Gogh/Strabismus) (*17, 18*), and Celsr (Fmi) (*19*), and two intracellular proteins, Dvl (Dsh) (*20, 21*) and Pk (*22*), which interact with Fzd and Vangl, respectively. Importantly, Fzd and Dvl, and Vangl and Pk are localized in an asymmetric manner at interfaces of adjacent cells, resulting in polarization of cell structures. One of the mysteries in PCP formation is, “How are individual cell polarities aligned?” One possible answer to this question is that a “global cue” controls the polarity of each cell. Since PCP can be directed by locally expressed Wnt in *Drosophila* and frog embryos (*23, 24*), a model has been proposed in which the polarity direction of each cell is determined depending on the direction of a Wnt concentration gradient (*7*). Consistently, some mutants of Wnt genes, as well as loss-of-function of components of a noncanonical Wnt signaling pathway, the Wnt/PCP pathway, exhibit abnormalities in PCP, or PCP-related phenotypes including convergent extension movements in vertebrates (*25–28*), although recent studies with *Drosophila* demonstrated that Wnt ligands are not required for PCP formation in the wing (*29, 30*). In the “global cue model”, Wnt ligands secreted from producing cells form a concentration gradient, along which each cell establishes polarity (*4, 7*). If we assume that this gradient covers the entire tissue in which PCP is formed, the gradient must be shallow, because such a tissue is usually much larger than the area of the signal-producing cells. However, in this situation, for each cell to sense the direction of the concentration gradient, cells must be sensitive enough to read slight concentration differences. On the other hand, if we assume that the sensitivity of the cells is high enough, the gradient must be very smooth and precise, otherwise the direction of PCPs will be disturbed. It remains unclear how Wnt is distributed in tissues where PCP is formed, but the smoothness of Wnt signaling gradient is disturbed by noise during morphogenesis along the A-P axis in zebrafish embryos (*31*). Therefore, although Wnt is actually required for PCP formation in many vertebrate tissues, it remains unclear whether the “global cue model” is adequate to explain the function of Wnt in PCP formation.

On the other hand, it has been shown in *Drosophila* and mice that interactions between neighboring cells are important for PCP formation (*32–34*). In contrast to the global cue model, these lines of evidence imply the importance of local cell-to-cell interaction for PCP formation. For instance, mouse chimeras consisting of PCP-sufficient cells mixed with cells that cannot form PCP indicate that core PCP components cannot be polarized cell autonomously. Rather, PCP-mediated contacts along single cell-cell interfaces is sufficient to polarize core PCP components (*34*). These results show that intercellular interactions among core PCP components are critical for PCP formation. However, the molecular mechanism underlying these intercellular interactions and the relationship between them and Wnt signal remains unclear.

To better understand the function of Wnt signaling in PCP formation, we tried to observe Wnt11 distribution in PCP formation. We have already succeeded in observing distribution of several Wnt ligands, including Wnt8, in embryonic tissues by generating antibodies for immunohistochemistry or by using Wnt ligands with fluorescent tags. Interestingly, Wnt8 was localized on cell surfaces in punctate patterns, suggesting that Wnt may not diffuse freely over wide areas, but that it accumulates locally on cell surfaces (*35, 36*). In this study, we created a Wnt11 antibody that can be used for immunostaining of embryos and we analyzed the distribution of endogenous Wnt11 using *Xenopus* embryos. In contrast to prevailing models, in the neural plate, where PCP is generated, Wnt11 was not distributed in a graded manner, but was polarized at cell boundaries in the direction of PCP instead. Surprisingly, even if Wnt is expressed ubiquitously, i.e., without any direction, core PCP components polarize autonomously, and at the same time, Wnt11 also polarizes, depending on core PCP components. This polarization can be generated by a positive amplification loop between Wnt11, Fzd7, and phosphorylated Vangl2 in a cis-configuration, and it can be transmitted to adjacent cells via asymmetric assembly of PCP components at cell boundaries. Thus, although Wnt11 and core PCP components do not initially have spatially biased distributions, their interactions and assembly spontaneously generate polarity in cells and tissues. Based on these results, we propose that Wnt11 does not regulate PCP in a top-down manner as a global cue, but rather can generate tissue polarity in a bottom-up manner by assembling locally with other proteins in cell membranes.

## Results

### Wnt11 is polarized in PCP formation

Transcripts for *Xenopus* Wnt11 (*wnt11b.L*) are strongly expressed in the posterior embryonic region during late gastrula (Stage (St.) 12) to early neurula (St. 14), which is the critical period for establishment of PCP in the neural plate (*37, 38*) (Fig. 1A and S1A). In addition, its expression was detectable in the anterior and middle regions of neural plate after St. 13 (Fig. 1A and S1A). To examine whether Wnt11 proteins are distributed in a graded manner from these sources, we performed whole-mount immunostaining using a newly generated anti-*Xenopus* Wnt11 antibody (for specificity, Fig. S1B and S1C). Wnt11 staining became intense in the neural plate during Stages 13-14, but it did not show a graded distribution in the posterior-to-anterior direction (Fig. 1B, 1C, S1D, and S1E). Surprisingly, precise imaging at higher magnification revealed that Wnt11 preferentially accumulated on cell boundaries in the medio-lateral direction of embryos (Fig. 1D and 1E). In addition, antisense morpholino oligo (MO)-mediated knockdown of a core PCP component gene, *vangl2*, reduced cell membrane-localized Wnt11 (Fig. 1F and 1G). These results suggest that Wnt11 localization and abundance on the cell membrane are under control of PCP in *Xenopus* embryos.

**Fig. 1.**
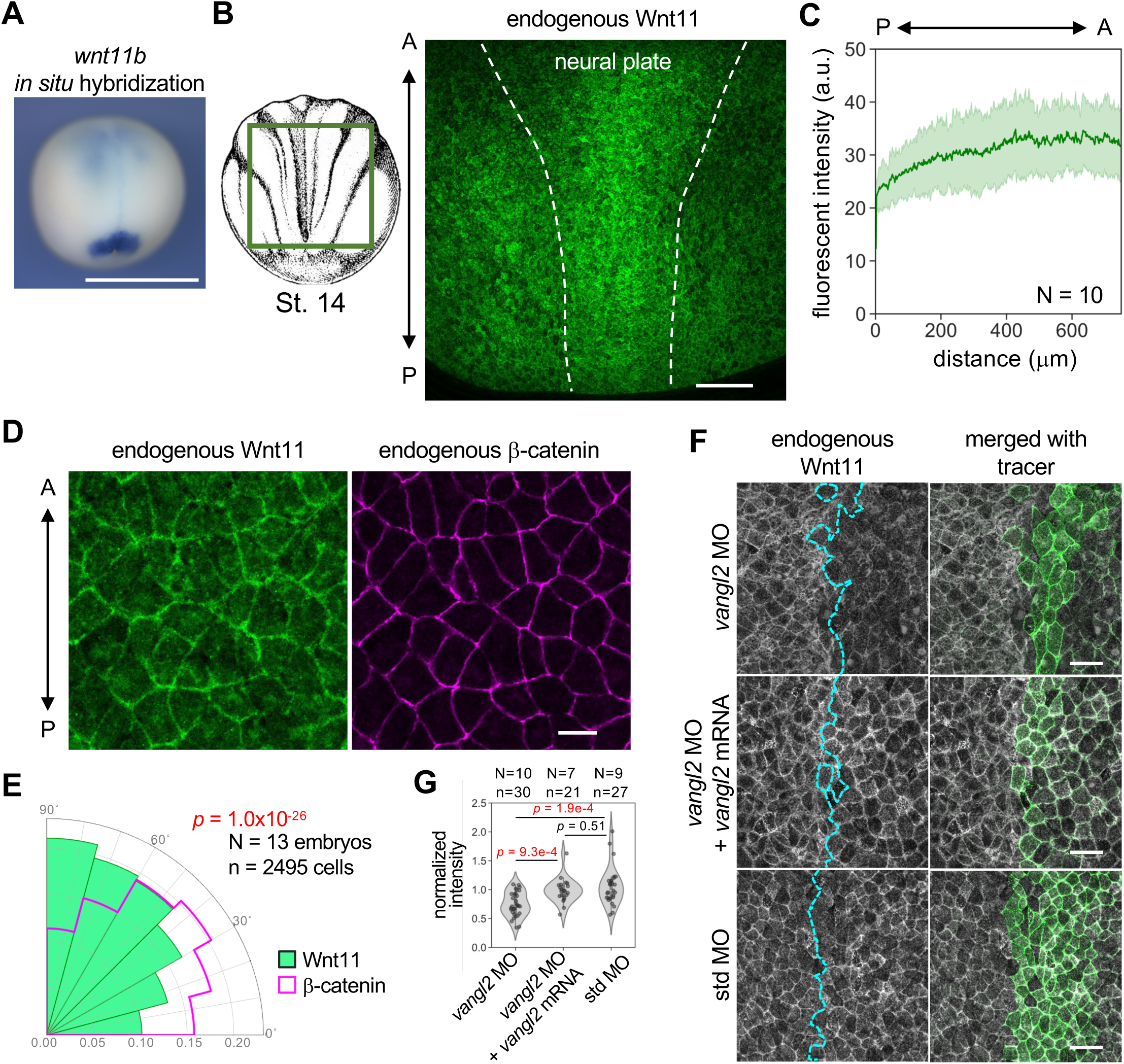
Wnt11 exhibits polarized localization in the neural plate in a PCP-dependent manner. **A, Expression pattern of *wnt11* transcripts in early neurula (St. 13).** Posterio-dorsal view. Expression of *wnt11b.L* mRNA was visualized by whole-mount *in situ* hybridization with an antisense probe. **B, Immunostaining of endogenous Wnt11 in *Xenopus* embryos (St. 14).** The observed area was as illustrated (left). The neural plate region is outlined with dashed lines. Wnt11 protein was concentrated in the neural plate, but in the neural plate, Wnt11 is widely distributed, even in the anterior part. The anterior-posterior axis is indicated. **C, Spatial profile of Wnt11 in the neural plate.** Wnt11 did not form a concentration gradient from posterior to anterior in the *Xenopus* neural plate. The light green area indicates one standard deviation. **D, Polarized localization of Wnt11.** The middle region of the neural plate is presented. β-catenin was also stained as a membrane marker without polarized localization. Notably, Wnt11 accumulates on cell boundaries in the medio-lateral direction (horizontal in the panel). **E, Quantification of polarized localization of Wnt11 in the neural plate.** Rose diagram (histogram) of polarity angle of Wnt11 (green) and β-catenin (magenta). Wnt11 exhibits significantly polarized localization toward 90° (corresponding to horizontal localization), compared to β-catenin. Kuiper two-sample test (a circular analog of the Kolmogorov–Smirnov test) test was used for statistical analysis. **F, Knockdown of *vangl2* with *vangl2* MO reduced membrane localization of Wnt11.** MOs with or without *vangl2* mRNA and membrane tracer (*mRuby2-krasCT* mRNA) were co-injected into the right dorsal blastomere of 4-cell embryos, targeting the future neural plate. Boundaries of tracer-negative and - positive areas are indicated with cyan dashed lines. **G, Quantifying reduction of Wnt11 membrane localization by *vangl2* knockdown.** Numbers of embryos (N) and numbers of ROIs (n) are as indicated. Numbers of embryos (N) and numbers of cells (n) are as indicated. Amounts of mRNAs/MOs (ng/embryo): *mRuby2-krasCT*, 0.067; *vangl2* MOs, 14; std MO, 14. Scale bars, 200 μm (A), 20 μm (D), 50 μm (F). Representative data from

In contrast, Wnt11 can induce PCP. For instance, PCP is ectopically established by Wnt11 in a reconstruction system using *Xenopus* embryos (*24*), referred to hereafter as reconstructed PCP (rPCP). In this system, Wnt11 is ectopically expressed in cells (source cells) adjacent to those expressing GFP-Pk3 and HA-Vangl2 (reporter cells) in the animal cap region (Fig. 2A), where PCP is rarely generated (*24*). To visualize Wnt11 in this system, we utilized mTagBFP2-tagged Wnt11 (Wnt11-BFP, W11B), which retained the ability to polarize GFP-Pk3 in rPCP (Fig. S2A). W11B exhibited polarized co-localization with GFP-Pk3 on reporter cells (Fig. 2B and S2A; This co-localization was apparent in a region in which GFP-Pk3 and HA-Vangl2 are sparsely expressed.) Furthermore, W11B polarization was also dependent on PCP because it was not observed in the absence of GFP-Pk3 (Fig. 2B). Thus, while Wnt11 induces PCP, PCP also localizes Wnt11 in rPCP.

**Fig. 2.**
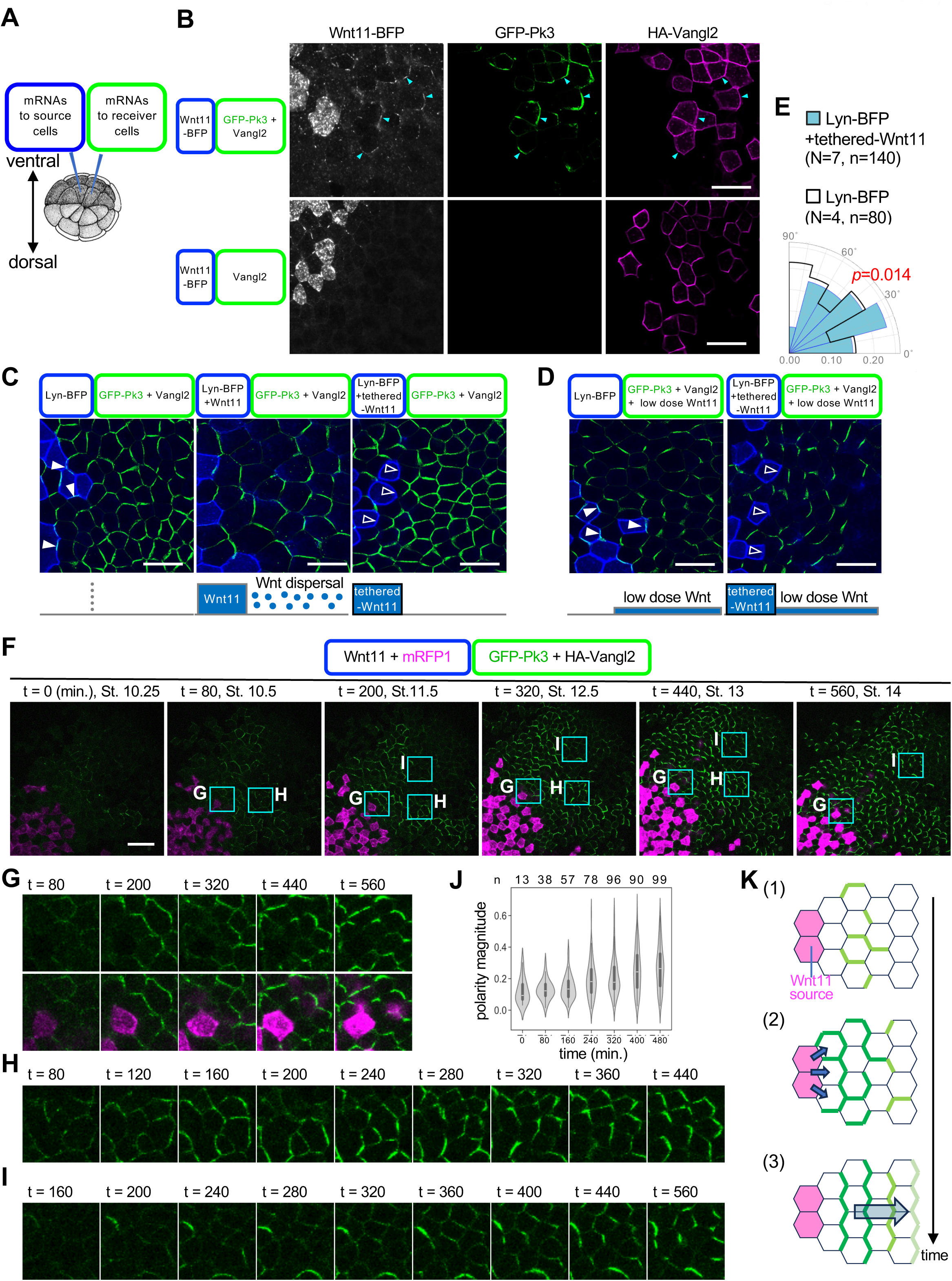
Directional and non-directional roles of Wnt11 for PCP and spatio-temporal profiling of Wnt-dependent polarization of GFP-Pk3. **A, Schematic illustration of mRNA injection for reconstructed PCP (rPCP).** Animal view of a *Xenopus* embryo at the 32-cell stage. **B, Localization of Wnt11 depends on formation of ectopic PCP.** Arrowheads indicate accumulations of Wnt11-BFP with GFP-Pk3 in a region in which GFP-Pk3 and HA-Vangl2 are sparsely expressed. Ectopic HA-Vangl2 without GFP-Pk3 is not sufficient for localization of Wnt11-BFP. **C, Tethered-Wnt11 cannot polarize GFP-Pk3 at long range, but does in adjacent cells.** Whereas diffusible Wnt11 can polarize GFP-Pk3 over a range of 5 cell diameters, tethered Wnt11 cannot. Note the presence (closed arrowheads) and absence (open arrowheads) of GFP-Pk3 at boundaries facing tracer- and tethered-Wnt11-expressing cells, respectively. **D, Ubiquitously expressed Wnt11 can polarize GFP-Pk3 without direction, and when combined with direction by tethered-Wnt11, polarization can be aligned long-range.** Note the presence (closed arrowheads) and absence (open arrowheads) of GFP-Pk3 at boundaries to tracer- and tethered-Wnt11-expressing cells, respectively. E, Quantification of polarity angles with or without direction by tethered-Wnt11. Numbers of embryos (N) and numbers of cells (n) are as indicated. **F-I, Time-lapse imaging of rPCP formation in the animal cap region.** mRFP1 was used as a tracer for Wnt11-expressing cells. (F) Overall, polarization appears to propagate from the vicinity of the Wnt11 source to distant regions. (G-I) Subregions adjacent (G, H) or distant (I) to the source are enlarged. **J, Quantification of polarity magnitude during time-lapse observation.** Polarization of GFP-Pk3 in the movie was quantified. Numbers of analyzed cells (n) are as indicated. K, Schematic illustrations of PCP formation in rPCP. mRNAs were injected into ventral animal blastomeres at the 32-cell stage, as illustrated. Amounts of mRNAs (pg/embryo): *wnt11-BFP*, 500; *lyn-BFP*, 50; *GFP-pk3*, 200 (B-E), 100 (F-J); *HA-vangl2*, 100 (B-E), 50 (F-J). *wnt11,* 500 (C, F-J), 50 (D); *tethered-wnt11*, 500. *mRFP1*, 500. Scale bars, 50μm.

### Wnt11 can induce PCP by a combination of directional and non-directional functions

To detail the reciprocal relationship between Wnt and PCP in rPCP, we focused on how Wnt regulates PCP formation. To examine whether dispersal of Wnt11 is required for polarization, unbound Wnt11 was replaced with membrane-tethered Wnt11-CD8-mCherry (denominated tethered-Wnt11). As predicted, this replacement abolished GFP-Pk3 polarization in reporter cells more than 2-cells away from cells expressing tethered-Wnt11 (Fig. 2C), indicating that diffusion-mediated dispersal of Wnt11 is required for long-range polarization. On the other hand, cell membrane localization of GFP-Pk3 was specifically abolished from the border adjacent to source cells (Fig. 2C and S2B), suggesting that Wnt11 tethered to cell membranes can polarize neighboring cells. Interestingly, when low-dose Wnt11 was additively and evenly expressed in reporter cells, long-range PCP was generated, directed toward tethered-Wnt11 expressing cells (Fig. 2D and 2E). Thus, although Wnt11 dispersal is required for PCP formation, a Wnt11 gradient in reporter cells is likely not essential for PCP formation.

Even in the absence of Wnt11 expression in source cells, additively and evenly expressed Wnt11 polarized GFP-Pk3 in reporter cells, although the direction of cell polarity was not consistent (Fig. 2D). Thus, even when acting without directional bias, Wnt11 can generate cell polarity, although it cannot determine the direction of polarity.

Based on these results, we speculate that Wnt11 contributes to PCP formation by at least two means. First, Wnt11 can induce polarization without directing polarity. In rPCP, this function appears to be contributed by Wnt11 molecules that diffuse ≥1 cell diameter from source cells. Second, Wnt11 from source cells polarizes adjacent cells, providing polarity direction toward Wnt-source cells. We refer these two functions as non-directional and directional polarization, respectively. These two functions likely represent characteristics of Wnt11-dependent formation of PCP such that together, they generate PCP.

### Live-imaging supports two distinct functions of Wnt11 in PCP formation

To see whether these two functions of Wnt11 are actually observed in formation of PCP, we followed dynamic processes in rPCP. By live imaging of this system from St. 10.25 to St. 14, we succeeded in observing the entire duration of PCP establishment (Movie S1 and Fig. 2F for cropped images). At the beginning, t = 0, GFP-Pk3 was very weakly localized on cell membranes. Around t = 80 min., which corresponds to St. 10.5, the first indications of biased localization of GFP-Pk3 were detected in cells some distance from, but not adjacent to Wnt11 source cells (Fig. 2G and 2H). At this time, the direction of GFP-Pk3 polarity was not persistent, fluctuating during a period of 20-40 min (Fig. 2H). On the other hand, in cells adjacent to Wnt11 source cells, polarization started around t = 200 min. (St. 11.5) (Fig. 2F and 2G). Interestingly, with some delay after this polarization in cells bordering Wnt11 source cells, the intensity of GFP-Pk3 in surrounding cells became more apparent. Its orientation became aligned, and its fluctuations decreased (Fig. 2H and 2J). Thus, once polarity was established in cells bordering the Wnt11 source, polarity in surrounding cells became more apparent and their polarity direction became stabilized. This observation can be explained by a combination of the two functions of Wnt11. The nondirectional function generates fluctuating polarity, and the directional function at the interface with the Wnt11 source gradually fixes the polarity of the non-directionally polarized cell population.

In addition to these processes, we also observed another aspect of GFP-Pk3 polarization. After the direction of PCP was once stabilized, polarization of GFP-Pk3 was propagated distally without undergoing non-directional polarization in areas distal to the cells shown above (Fig. 2I). This suggests that PCP can propagate distally once stable PCP is established (Fig. 2K).

### Polarity is generated via reciprocal interaction between Wnt11 and core PCP components

In non-directional polarization, one of the main questions is how polarization of molecules is generated without directional cues. To address this question, we examined spatial patterns of Wnt11 and core PCP components under conditions in which Wnt11 is not provided by distant source cells, but is expressed only in reporter cells. To visualize Wnt11, Vangl2, and Pk3, we utilized W11B, monomeric EGFP-tagged Vangl2 (GV2) and mRuby2-tagged Pk3 (RP3), respectively. We also examined the spatial pattern of endogenous Fzd7 by immunostaining. Because stability of endogenous Fzd7 was increased simply by overexpressing Vangl2 in the same cells (Fig. S3D and S3E and S7A and S7B), we expressed GV2 in all experiments. First, we confirmed that addition of these tags did not perturb cell polarization because co-expression of W11B, GV2 and RP3 can polarize all these components, as well as endogenous Fzd7 (Fig. 3A first row). As predicted, in the absence of W11B, polarity of GV2, Pk3 and Fzd7 disappeared, showing that their polarity is dependent on Wnt11 (Fig. 3A second row). On the other hand, Wnt11 reduced mean Pk3 levels in a dose-dependent manner, but at the same time, an intermediate dose of Wnt11 accumulated locally and polarized Pk3, indicating that appropriate levels of Wnt11 expression are important for polarization (Fig. S4A). In the absence of RP3, polarity of GV2 almost disappeared, but Wnt11 and endogenous Fzd7 accumulated together with slight polarity (Fig. 3A third and fourth rows), suggesting that Pk3 enhances Wnt11-dependent polarity. These results suggest that cell polarity can be established without directional cues, but only through reciprocal interaction between Wnt11 and core PCP components, including Vangl, Pk, and Fzd.

**Fig. 3.**
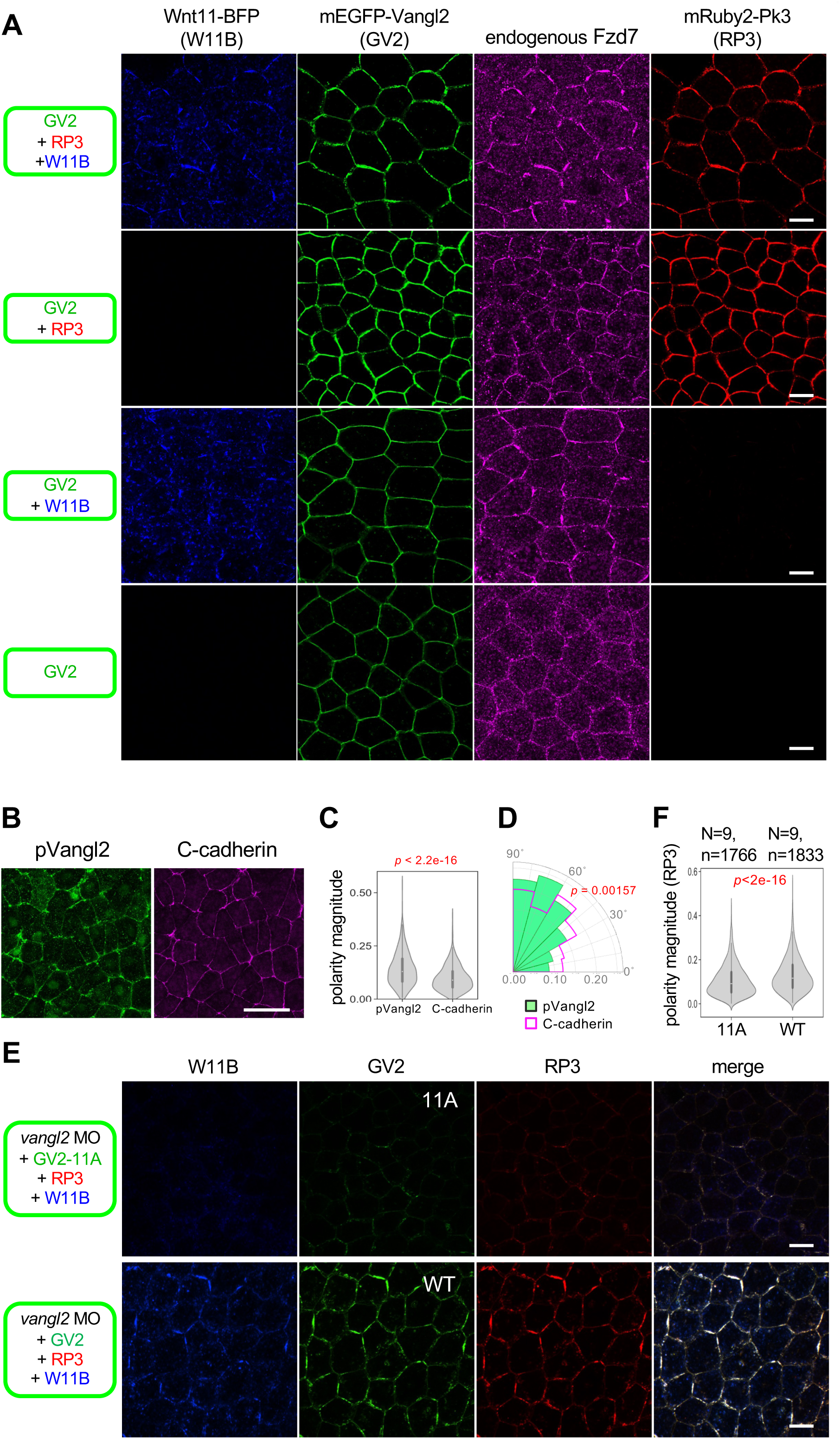
Localization of core PCP components in non-directional polarization by ubiquitous Wnt11. **A, Localization of core PCP components in the absence or presence of Wnt11.** Co-expression of Wnt11-BFP (W11B) polarized membrane localization of GV2, RP3, and endogenous Fzd7. In addition, W11B co-localizes with these core PCP components. In the absence of W11B, membrane localization of GV2 was increased by co-expressed RP3, but its membrane localization was uniform. **B-D, pVangl2 is polarized in the neural plate.** (B) pVangl2 staining. C-cadherin was used as a membrane marker without polarization. (C) Polarity magnitude of pVangl2 is significantly higher than that of C-Cadherin. (D) The polarity angle of pVangl2 and C-Cadherin. Data of 5 embryos, 699 cells are presented (C, D). Statistical analyses were performed with the Kuiper two-sample test for polarity angle (D). Polarity magnitude and polarity angle were quantified with the PCA method using QuantifyPolarity2.0 (*43*). **E, F, Phosphorylation of Vangl2 is required for polarized localization of core PCP components and Wnt11.** (E) Substitution of GV2 with a phospho-deficient form (GV2-11A) leads to reduction of all observed components. (F) Quantification of PCA magnitude indicates a significant loss of polarization with GV2-11A, compared with wild-type GV2. Numbers of analyzed embryos (N) and cells (n) were as indicated. mRNAs and *vangl2* MO were injected into the animal pole region of a ventral blastomere at the 4-cell stage, as indicated. Amounts of mRNAs/MOs (ng/embryo): *mEGFP-vangl2* (WT and 11A), 0.10; *mRuby2-pk3*, 0.20; *wnt11-BFP*, 1.0. *vangl2* MO, 21. Scale bars, 20 μm (A, E), 50 μm (B).

### Co-existent phosphorylated and non-phosphorylated states of Vangl2 are essential for polarization

To better understand the molecular mechanism of non-directional function of Wnt11, we focused on downstream events of Wnt11 signaling. Interestingly, Vangl phosphorylation at the N-terminal region, which leads to Vangl2 stabilization by avoiding ubiquitination (*39*) and is essential for PCP (*27, 40–42*), is induced by Wnt5a in the context of PCP formation in mammalian development (*27, 41*). Immunostaining of pVangl2 by anti-pVangl2 antibody (see Fig. S5A-E for specificity) showed pVangl2 staining preferentially accumulated at cell boundaries along the medio-lateral direction of embryos (Fig. 3B), highly correlated with endogenous Wnt11 staining (Fig. S5F-H) and overexpressing W11B (Fig. S5I). Furthermore, overexpression of Wnt11 increased pVangl2 staining in the animal cap region (Fig. S5A-C).

These results suggest that Vangl2 phosphorylation is involved in Wnt11-induced PCP formation in *Xenopus* embryos.

To examine whether Vangl2 phosphorylation is required for the non-directional function of Wnt11, we replaced Vangl2 with mEGFP-tagged phospho-deficient Vangl2 (GV2-11A) in *Xenopus* embryos, under conditions in which endogenous Vangl2 was depleted by *vangl2*-specific MO. GV2-11A, but not wild-type GV2 (both are MO-resistant) impaired Pk3 polarization (Fig. 3E and 3F), showing that phosphorylation of Vangl2 is crucial for Wnt11-dependent non-directional polarization.

Next, to test the sufficiency of Vangl2 phosphorylation for Wnt11-induced polarization, we examined whether a phospho-mimetic form of Vangl2, HA-Vangl2-11D (HV2-11D) can promote RP3 polarization in the absence of Wnt11 under Vangl2-depleted conditions with vangl2 MO. As a control, mEGFP-tagged phospho-deficient Vangl2 (GV2-11A) was also used. Quantification with polarity magnitude (*43*) showed that neither HV2-11D nor GV2-11A can induce RP3 polarization (Fig. 4A and 4B and 4H). Notably, co-expression of GV2-11A and HV2-11D led to polarization of RP3 (Fig. 4D and 4H). In this condition, GV2-11A and HV2-11D were also polarized in a Pk3-dependent manner (Fig. 4C and 4D). Thus, concurrent phosphorylated and non-phosphorylated states of Vangl2 appear important for polarization.

**Fig. 4.**
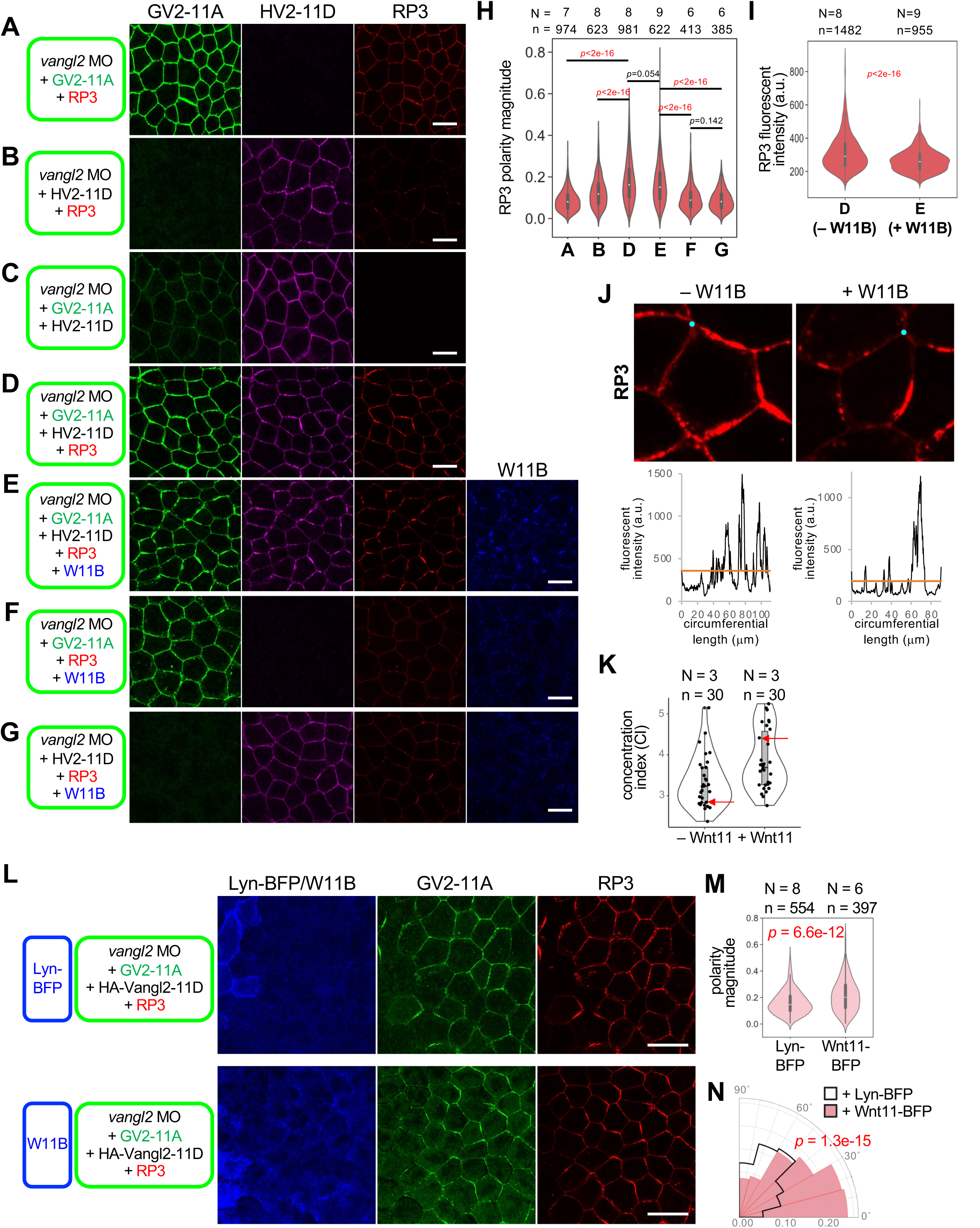
Reconstitution of cell polarity based on phosphorylated and unphosphorylated states of Vangl2. **A-G, A combination of Vangl2-11A, -11D and Pk3 is sufficient for self-polarization with or without Wnt11.** In all conditions, endogenous Vangl2 was knocked down with *vangl2* MO. mRNAs and *vangl2* MO were injected into the animal pole region of a ventral blastomere at the 4-cell stage, as indicated. **H, Quantification of polarity magnitude of mRuby2-Pk3 in reconstitution of polarity.** Polarity magnitude was quantified with the PCA method using QuantifyPolarity2.0 (*43*). **I, Quantification of RP3 on cell membranes.** Conditions of – Wnt11 (D) and + Wnt (E) were compared. **J,K, Co-expression of Wnt11 concentrates Pk3 around cell circumference.** (J) Representative images of RP3 with or without Wnt11. Graphs show circumferential plots of RP3 (counterclockwise, starting from points indicated with cyan dots), with mean values of RP3 intensity of each cell (orange lines). (K) Concentration index (CI). CIs were calculated as follows: CI = [circumferential length of the cell]/[length with signal intensity above the mean circumferential intensity of the cell]. Arrows indicate examples shown in **J**. Co-expression of Wnt11 significantly increased CI. **L-M, Wnt11 overexpressed in an adjacent region can further polarize and direct Pk3.** mRNAs and *vangl2* MO were injected into the animal pole region of ventral blastomeres at the 4-cell stage as indicated, as in rPCP. Statistical analyses were performed with the Kuiper two-sample test for polarity angle (N). Numbers of embryos (N) and numbers of cells (n) are as indicated. Amounts of mRNAs/MOs (ng/embryo): *mEGFP-vangl2-11A*, 0.050; *HA-vangl2-11D*, 0.025; *mRuby2-pk3*, 0.10; *wnt11-BFP*, 0.50; *vangl2* MO, 21. Scale bars, 20 μm (A-G), 50 μm (F).

Unexpectedly, when W11B was co-expressed with GV2-11A, HV2-11D, and RP3 (Fig. 4E), overall, RP3 levels were reduced (Fig. 4I), but it became more concentrated on cell membranes (Fig. 4J and 4K), although polarity magnitude was not significantly different (Fig. 4H). This effect is not due to phosphorylation of Vangl2, because neither GV2-11A nor -11D can be further phosphorylated by W11B. Thus, in addition to effects via Vangl2 phosphorylation, Wnt11 affects localization of core PCP components independently of this phosphorylation reaction.

Finally, we confirmed that co-existence of phosphorylated and non-phosphorylated Vangl2 is not only essential for non-directional polarization, but also for establishment of directional PCP, as shown in rPCP (*24*) (Fig. 2). When W11B was expressed in cells adjacent to those expressing GV2-11A, HV2-11D and RP3 (Fig. 4L), the polarity angle of RP3 was directed toward the Wnt11 source, and the polarity magnitude of RP3 was also significantly increased (Fig. 4M and 4N). Together, these results suggest that both Wnt11-induced phosphorylated and non-phosphorylated states of Vangl2 appear important for establishment PCP in *Xenopus* embryos. At the same time, these results also suggest that Wnt11 contributes to PCP formation independently of this phosphorylation reaction.

### Phosphorylated Vangl2 is polarized on the side opposite Pk3

Vangl2 phosphorylation is dependent on Fzd, but is inhibited by Pk (*40, 44*). Furthermore, this phosphorylation inhibits interaction between Vangl2 and Pk3 in *Xenopus* embryos (*44*). Although Vangl localizes on Pk-rich cell membranes in polarized cells, these findings imply that pVangl2 does not. Thus, we minutely examined localization of pVangl2 at the cell boundary using super-resolution stimulated emission depletion (STED) microscopy, which was able to resolve localization of membrane tracers expressed in adjacent cells (Fig. S6A-C), to analyze non-directional polarization (see Fig. 2D and S4).

STED observations showed that Wnt11 segregates Pk3 and Fzd7 staining (Fig. 5A and 5B), corresponding to their localization on opposite sides at cell boundaries of polarized cells. Notably, in this condition, pVangl2 staining was segregated from GFP-Pk3 (Fig. 5C, 5D, and S6D), suggesting that Vangl2 phosphorylation specifically occurs on the Fzd-side in polarized cells (Fig. 5E). Consistent with this hypothesis, at the cell membrane facing Wnt11-expressing cells, pVangl2 was clearly observed, along with disappearance of GFP-Pk3 (Fig. 5F).

**Fig. 5.**
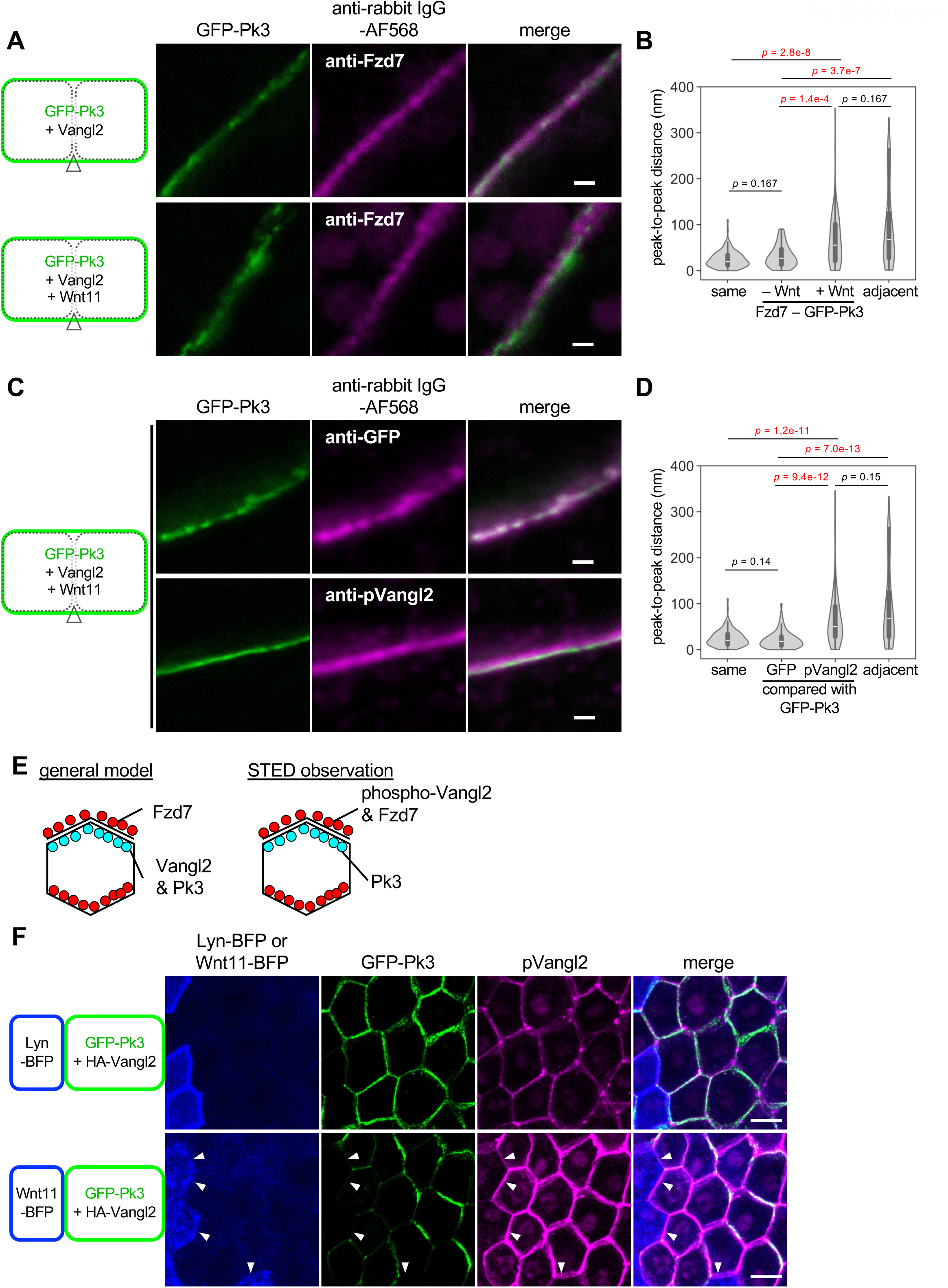
Localization of phosphorylated Vangl2 in polarized cells analyzed by super-resolution microscopy (STED). **A, B, STED observations of GFP-Pk3 and endogenous Fzd7 with or without Wnt11.** (A) Conditions with or without Wnt11 (thus polarized or not polarized, respectively) were compared. (B) Peak-to-peak distance (PPD) between Fzd7 and GFP-Pk3. Data from 3 embryos, 15 cell boundaries, 67 lines (–Wnt) and 4 embryos, 18 cell boundaries, 76 lines (+Wnt) are presented. For “same” and “adjacent”, same data as in Fig. S6C are presented. **C, D, STED observations of GFP-Pk3 and phosphorylated Vangl2.** (C) GFP-Pk3 and anti-GFP staining or anti-pVangl2 staining. Cells polarized by overexpression of Wnt11 were analyzed. Anti-GFP staining was used as a control that overlapped with GFP fluorescence. (D) PPD between GFP-Pk3 and anti-pVangl2 staining. Data from 3 embryos, 14 cell boundaries, 74 lines (anti-GFP) and 3 embryos, 16 cell boundaries, 90 lines (anti-pVangl2) are presented. For “same” and “adjacent”, same data as in C are presented. **E, Localization of core PCP components based on STED microscopy.** In general models, Vangl2 localizes on the same side as Pk3. Our STED analyses suggest that phosphorylated Vangl2 localizes on the opposite side from Pk3, possibly the same side as Fzd7. Because Pk3 preferentially binds to nonphospho-Vangl2 (*44*), we speculate that nonphospho-Vangl2 is on the same side as Pk3 in a polarized cell. **F, pVangl2 localized to cell membranes where GFP-Pk3 was reduced by Wnt11 in rPCP.** mRNAs were injected as illustrated. pVangl2 was weak at cell boundaries adjacent to Lyn-BFP-expressing cells, but strong at boundaries adjacent to Wnt11-BFP-expressing cells. All STED pictures in Fig. 4 and S6 are shown in the same condition for brightness/contrast. mRNAs were injected into animal pole regions of ventral blastomeres at the 4-cell stage, as indicated. Amounts of mRNAs (ng/embryo): *GFP-pk3*, 0.20; *vangl2*, 0.10; *HA-vangl2*, 0.10; *wnt11*, 0.25; *wnt11-BFP*, 0.50; *lyn-BFP*, 0.050. Scale bars, 500 nm (A, C), 20 μm (F).

### Wnt11 induces assembly of phosphorylated Vangl2 and Fzd7 on the same cell membrane

Our analysis with STED microscopy suggests that Wnt11 induces assembly of pVangl2 with Fzd7 on the same membrane. Since overexpressed Vangl2 increased Fzd7 amounts on the same membrane (Fig. S7C-S7F and S8), we asked whether and how Wnt11 influences Fzd7 via phosphorylation of Vangl2 on the same membranes. For this purpose, mRuby2-tagged Fzd7 (Fzd7-mRuby2, F7R) was expressed with GV2, GV2-11A, or a phospho-mimetic form, GV2-11D, in a group of cells. Their outer boundaries facing neighboring cells that did not express GV2 or F7R were analyzed in conditions with or without W11B. With GV2-11A and -11D, the effect of endogenous Vangl2 was depleted with a *vangl2*-pecific MO (Fig. 6D). While GV2 and F7R exhibit uniform localization along the outer boundaries (Fig. 6A), they assembled with W11B when W11B was expressed in adjacent cells (Fig. 6A and 6B). Thus, Vangl2 and Fzd7 can co-accumulate on the same membrane while interacting with Wnt11 (Fig. 6C, the degree of assembly of Vangl2, Fzd7, and W11B is evaluated with the concentration index (CI)). To distinguish these assemblies on the same membranes from those bridging opposing membranes, the former are referred to hereafter as cis-assemblies and the latter as trans-assemblies.

**Fig. 6.**
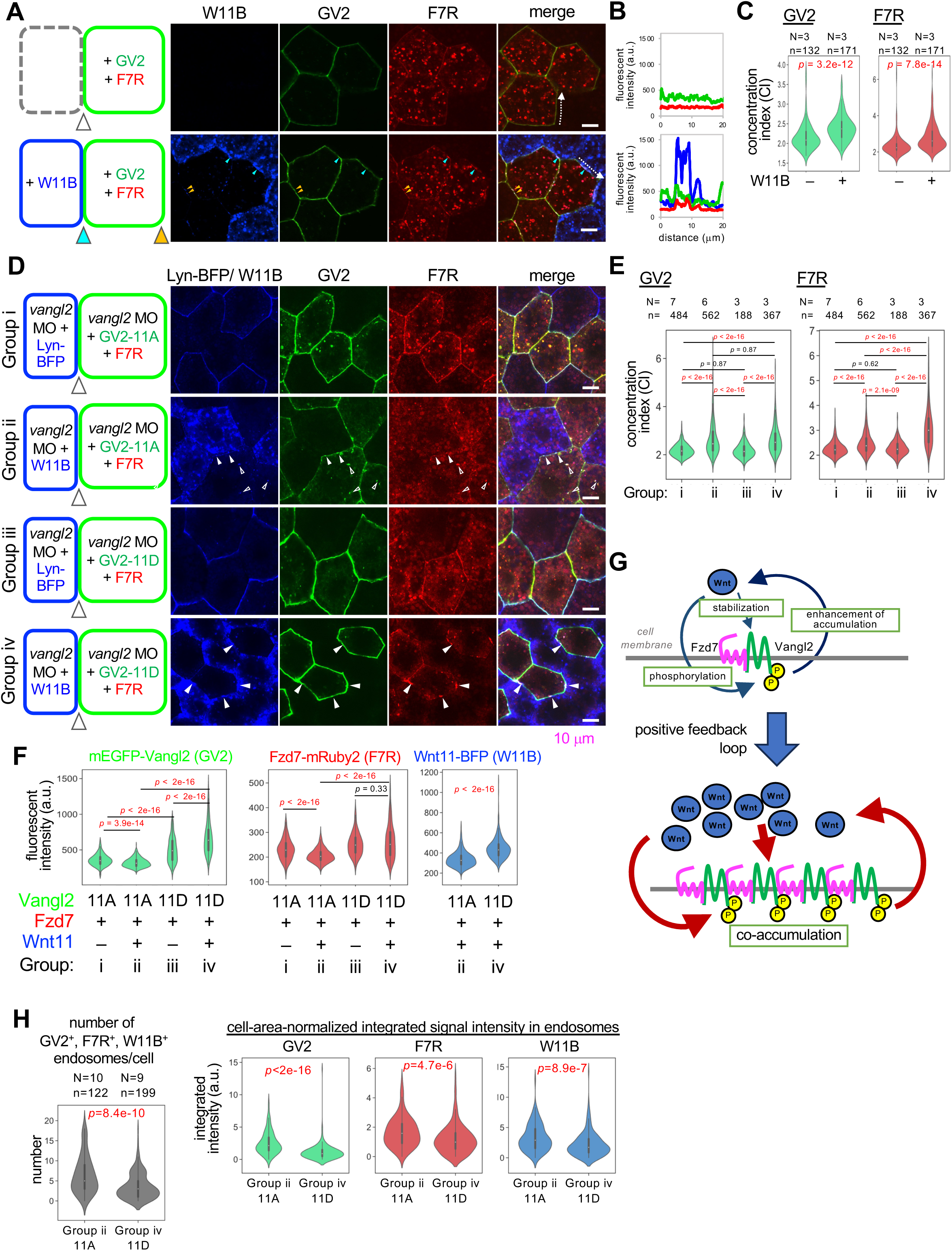
Cis-assembly of Vangl2 and Fzd7 induced by Wnt11, involving phosphorylation states of Vangl2. **A-C, Wnt11 induces cis-assembly of Vangl2 and Fz7.** (A) Cis-assembly is frequently observed on outer boundaries facing W11B-expressing cells (cyan arrowheads), but also on other outer boundaries (orange arrowheads). (B) Intensity profiles of each signal along the outer cell boundaries shown by dotted arrows (blue, W11B; green, GV2; red, F7R). (C) Wnt11 increased concentration indices of GV2 (left) and F7R (right). **D-F Effects of Vangl2 phosphorylation states on cis-assembly with Fzd7.** (D) Vesicles containing GV2-11A, F7R, and W11B are frequently observed inside cells (open arrowheads), indicating endosomal internalization. Cis-assembly of GV2-11D and F7R are frequently observed on cell membranes (closed arrowheads). (E) CIs of GV2 (left) and F7R (right) in each experimental group. (F) Quantification of tagged proteins. Outer cell boundaries of GV2- and F7R-expressing cells are quantified (arrowheads in each schematic image in (D)). Numbers of embryos (N) and numbers of cell boundaries (n): Group i, N = 7, n = 485; Group ii, N = 6, n = 562; Group iii, N = 3, n = 188; Group iv, N = 3, n = 367. G, Schematic illustration of co-assembly of phosphorylated Vangl2-Fzd7 cis-complexes by Wnt11 with positive feedback loops. **H, Phospho-deficient Vangl2 was highly internalized compared with phospho-mimetic Vangl2 in the presence of Wnt11.** Note that W11B vesicles detected in GV2- and F7R-co-expressing cells must be internalized from the extracellular space. Numbers of embryos (N) and numbers of cells (n) are as indicated (C, E, H). mRNAs and *vangl2* MO were injected into the animal pole region of ventral blastomeres at the 4-cell stage, as indicated. Amounts of mRNAs/MOs (ng/embryo): *mEGFP-vangl2*, 0.20; *mEGFP-vangl2-11A*, 0.10; *mEGFP-vangl2-11D*, 0.10; *fzd7-mRuby2*, 0.20 (A-C), 0.050 (D), 0.10 (E,F,H); *wnt11-BFP*, 1.0 (A-C), 0.50 (D-F, H); *lyn-BFP*, 0.050; *vangl2* MO, 42 (D), 28 (E,F,H). Scale bars, 10 μm.

Assembly of Vangl2 and Fzd7 with Wnt11was also observed even with GV2-11A (Fig. 6D and 6E, Group i vs ii), suggesting that Wnt11 can recruit Vangl2 and Fzd7 even when Vangl2 is not phosphorylated. On the other hand, this assembly was more continuous and intense with GV2-11D (Fig. S9B and S9D), and GV2-11D concentrated F7R more than did GV2-11A (Fig. 6D, E, Group ii vs iv). Thus, Vangl2 phosphorylation further promotes Wnt11-dependent cis-assemblies. Since this phosphorylation is dependent on Wnt/Fzd signaling (*27, 44*), our results suggest that Vangl2 phosphorylation and Wnt11/Fzd7 signaling promote one another, forming a positive-feedback loop that enhances assembly of Vangl2, Fzd7 and Wnt11 (Fig. 6G). In addition, the abundance of GV2-11D on outer boundaries was higher than that of GV2-11A, probably due to differences of transport efficiency from the ER to the cell membrane, as previously reported (*39*), and those of F7R and W11B were also higher with Vangl2 phosphorylation (Fig. 6F, Group i vs iii and Group ii vs iv, Fig. S7C-F, S8). Thus, Vangl2 phosphorylation may increase the abundance of Vangl2 itself, as well as of Fzd7 and Wnt11 at the outer border (Fig. 6G).

Abundance of Vangl2, Fzd7, and W11B on outer boundaries appears to show additional functions of Wnt11 in cis-assembly. GV2-11A was decreased by W11B (Fig. 6F, Group i vs ii), whereas GV2-11D was increased (Fig. 6F, Group iii vs iv). Wnt11 also reduced the amount of F7R on the same membranes as GV2-11A (Fig. 6F, Group i vs ii). Given that GV2-11A and -11D may not be further phosphorylated, this result suggests that some Wnt11 function other than inducing Vangl2 phosphorylation regulates stability of Vangl2 and Fzd7, depending on phosphorylation states of Vangl2. Thus, Wnt11 not only increases phosphorylation of Vangl2, but also further stabilizes phosphorylated Vangl2, thereby promoting assembly of Wnt11, Fzd7, and Vangl2.

Interestingly, numbers of intracellular puncta positive for GV2, F7R, and Wnt11 were significantly higher with GV2-11A than with GV2-11D (Fig. 6H and S9E). Considering that intracellular Wnt11 was only taken up from outside the cell, these results suggest that non-phosphorylated Vangl2 tends to be removed with Fzd7 and Wnt11 from the cell membrane via endocytosis (Fig. S9I). Furthermore, when RP3 was expressed with W11B and GV2-11A, RP3 on cell membranes was reduced with GV2-11A by W11B, and their endosome-like puncta were increased by W11B (Fig. S9F, same condition as Fig. 4A and 4F, and Fig. S9H). In addition, in embryos in which non-directional polarization was observed, GFP-Pk3 was often found in endosomes that were negative for pVangl2 staining (Fig. S9G). Thus, it is plausible that Pk3, non-phosphorylated Vangl2, and Fzd7 are coupled so as to be internalized into cells upon Wnt11 stimulation (Fig. S9I). This Wnt11/Fzd7-mediated internalization machinery may result in exclusion of Pk3 and non-phosphorylated Vangl2 from membrane regions enriched in Wnt11 and pVangl2.

### A combination of regulatory loops leads to asymmetric arrangement of core PCP components

Because our STED analysis suggests that Pk3 is mostly localized on the opposite side of pVangl2, we examined whether phosphorylation of Vangl2 impairs stability of Pk3 on the same membrane. As predicted, GV2-11A significantly increased mRuby2-Pk3 (RP3), whereas GV2-11D did not (Fig. 7A and 7B right). On the other hand, RP3 increased GV2-11A, but decreased GV2-11D (Fig. 7A and 7B left). Thus, Pk3 and non-phospho-Vangl2 appear to stabilize each other on the opposite side from pVangl2 and Fzd7 (Fig. 7C).

**Fig. 7.**
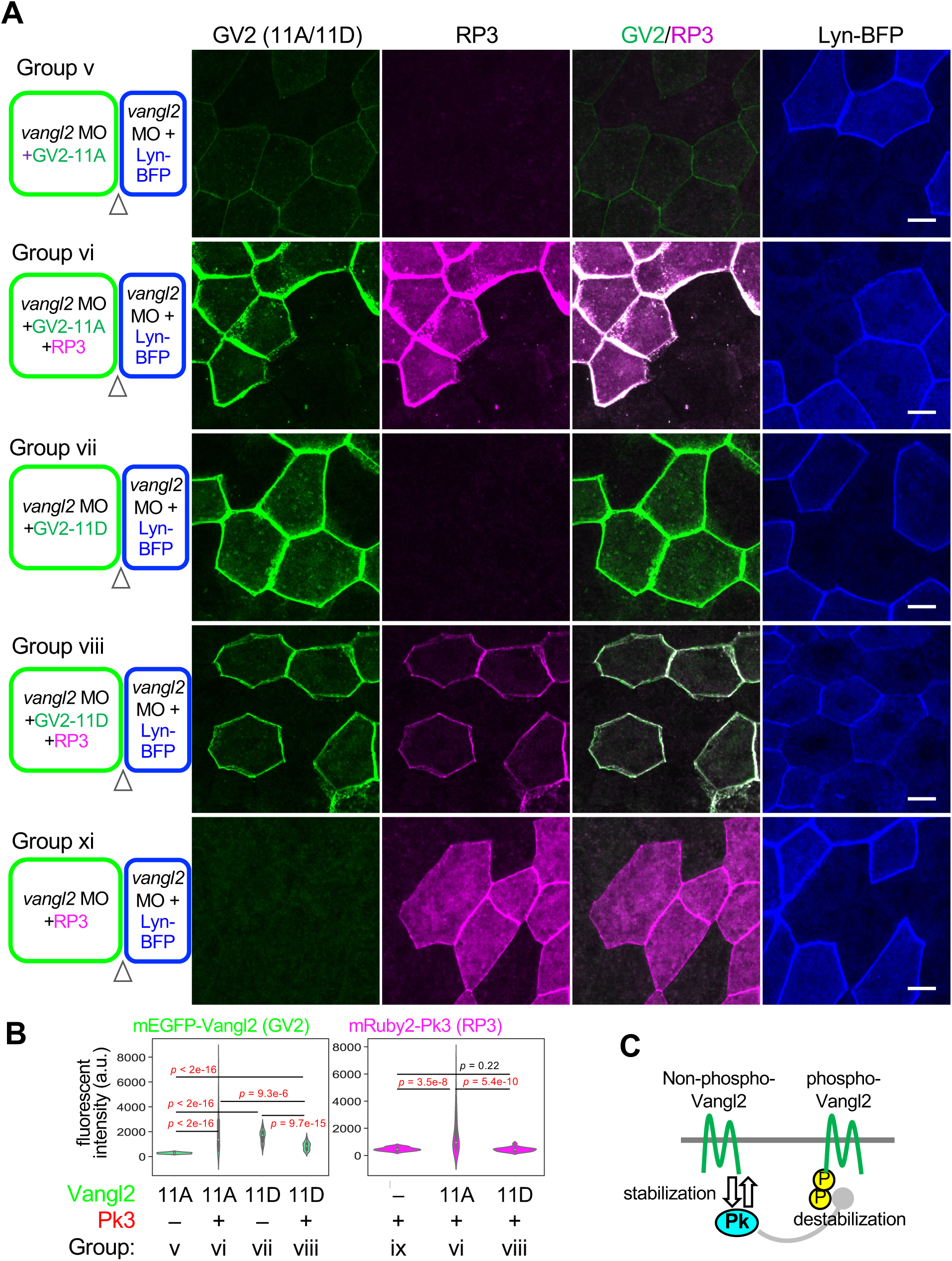
Reciprocal stabilization of Pk3 and non-phosphorylated states of Vangl2. **A, B, Effects of Vangl2 phosphorylation states with Pk3 on the same membrane.** (A) Five experimental groups (Group v – xi) were examined as indicated. (B) Outer cell boundaries of GV2 and F7R expressing cells, where both are on the same membrane, are quantified (arrowheads in schematic illustrations) for amounts of tagged proteins. Numbers of embryos (N) and numbers of cell boundaries (n): Group v, N = 7, n = 112; Group vi, N = 4, n = 55; Group vii, N = 5, n = 94; Group viii, N = 7, n = 55; Group ix, N = 5, n = 46. **C, Schematic illustration of stabilization/destabilization of Vangl2-Pk3 cis-complexes switched with phosphorylation states of Vangl2.** mRNAs and *vangl2* MO were injected into animal pole regions of ventral blastomeres at the 4-cell stage, as indicated. Amounts of mRNAs/MOs (ng/embryo): *mEGFP-vangl2-11A*, 0.10; *mEGFP-vangl2-11D*, 0.10; *mRuby2-pk3*, 0.20; *lyn-BFP*, 0.050; *vangl2* MO, 42. Scale bars, 10 μm.

In PCP, core PCP components are asymmetrically localized across cell boundaries, i.e., asymmetry with Pk and Fzd (see Fig. 5E) and are likely to accumulate in a trans-configuration, bridging adjacent cells (*32–34, 45*). We found that overexpressed W11B promoted trans-assembly of GV2, F7R, and W11B (Fig. 8A, 8B, and S10A). When endogenous Vangl2 was knocked down on the side of F7R, GV2-F7R trans-assembly was reduced (Fig. 8C, 8D, and S10B), indicating that Vangl2 on the Fzd-side is prerequisite for Vangl2-Fzd7 trans-assembly. To examine whether phosphorylation of Vangl2, which is crucial for cis-assembly with Fzd7, is also important for trans-assembly, non-tagged Vangl2-11A or -11D was expressed on the same side as F7R under conditions in which GV2-11A was expressed on the opposite side. Cis-Vangl2-11D significantly increased the level of GV2-11A and F7R, compared to controls without exogenous Vangl2 and Vangl2-11A, suggesting that cis-assembly of phosphorylated-Vangl2 and Fzd7 increases the level of non-phosphorylated Vangl2 on the opposite side (Fig. 8E, 8F and S10C).

**Fig. 8.**
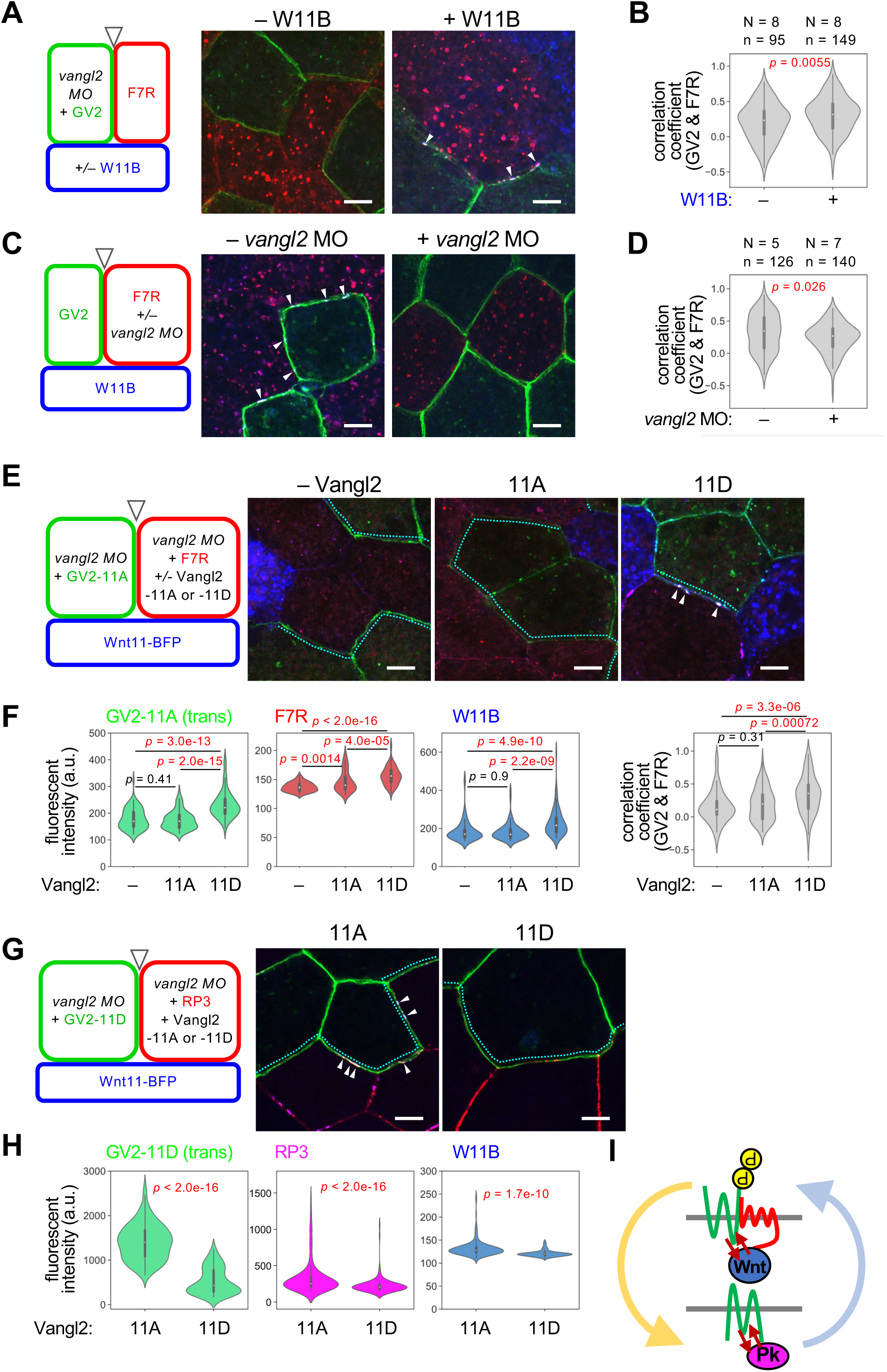
Trans-assembly of core PCP components by Wnt11. **A, B, Wnt11 induces trans-assembly of Vangl2 and Fzd7.** Cell boundaries between GV2-expressing and F7R-expressing cells were observed. (A) Overexpression of W11B induced co-accumulation of GV2, F7R and W11B (arrowheads), indicating trans-assemblies of Vangl2 and Fzd7 containing Wnt11. (B) Correlation coefficients between GV2 and F7R on opposing membranes. **C, D, Vangl2 is required on the same membranes as Fzd7 for trans-assembly of Vangl2 and Fzd7.** Boundaries between GV2-expressing and F7R-expressing cells were observed (arrowheads in schematic illustrations). (C) Co-accumulation of GV2 and F7R (arrowheads) was inhibited when *vangl2* MO was injected into the F7R side. (D) Correlation coefficients of GV2 and F7R on opposing membranes. **E, F, Vangl2-11D on the same membrane as Fzd7 can specifically increase Vangl2-11A on the opposite side.** Boundaries between GV2-expressing and F7R-expressing cells were quantified (cyan dotted lines). Not only protein amounts of GV2-11A, but also correlation coefficients between trans-GV2-11A and F7R were significantly higher with cis-Vangl2-11D. Numbers of embryos (N) and numbers of cell boundaries (n): cis-Vangl2 (–), N = 4, n = 97; 11A, N = 4, n = 81; 11D, N = 6, n = 94. **G, H, Vangl2-11A on the same membrane as Pk3 can specifically increase Vangl2-11D on the opposite side.** Boundaries between GV2-expressing and RP3-expressing cells were quantified (cyan dotted lines). Numbers of embryos (N) and numbers of cell boundaries (n): 11A, N = 8, n = 177; 11D, N = 4, n = 110. I, Schematic illustration of reciprocal stabilization of opposing phospho- and non-phospho-Vangl2, corresponding to Fzd7- and Pk3-sides, respectively. Numbers of embryos (N) and numbers of cell boundaries (n) are as indicated (B, D). mRNAs for GV2, GV2-11A/D, F7R, Vangl2-11A/D and *vangl2* MO were injected into the animal pole region of ventral blastomeres, and mRNA for W11B was injected into the animal pole region of dorsal blastomeres at the 4-cell stage, as indicated. Amounts of mRNAs/MOs (ng/embryo): *mEGFP-vangl2*, 0.10 (A, B), 0.20 (C, D); *mEGFP-vangl2-11A*, 0.10; *mEGFP-vangl2-11D*, 0.20; *fzd7-mRuby2*, 0.10 (A, B), 0.20 (C, D), 0.050 (E); mRuby2-Pk3, 0.20; *wnt11-BFP*, 1.0 (A, B, E, F), 2.0 (C, D); *vangl2-11A/D*, 0.10 (E), 0.050 (F); *vangl2* MO, 11 (A, B), 21 (C, D), 57 (E), 42 (F). Scale bars, 10 μm.

In turn, we examined whether Vangl2-Pk3 cis-interaction affects the level of phosphorylated-Vangl2 on the opposite side. In this case, non-tagged Vangl2-11A or -11D was expressed on the same side as RP3 under conditions in which GV2-11D was expressed on the opposite side. We found that Vangl2-11A increased the GV2-11D level more than Vangl2-11D did (Fig. 8G, 8H and S10D). These results indicate that the two types of cis-interactions either involving Fzd7 or Pk3 form on different sides of the cell boundary and stabilize formation of one another through trans-interaction. Given that Wnt11/Fzd7/pVangl2-mediated cis-assembly is amplified by a positive feedback loop, this amplification may spill over into the trans-interaction and even to Vangl2-Pk3 cis-interaction, leading to coordinated amplification of symmetry breaking across the cell boundary (Fig. 8I).

## Discussion

It is still debatable whether the concentration gradient of Wnt ligands serves as a directional cue for PCP. For instance, Vangl2 phosphorylation, which is induced by Wnt5a through Ror2, exhibits a gradient along the proximo-distal axis in the limb bud (*27*). Although this suggests that a Wnt5a signaling gradient is generated along the proximo-distal axis of the limb bud, it remains unclear whether Wnt ligands themselves simply form concentration gradients during PCP formation. Furthermore, it is also puzzling how each cell in a gradient senses the direction of the surrounding Wnt gradient and establishes its own polarity.

In this study, we found that Wnt11 does not form simple gradients as previously predicted (Fig. 1A-C, S1D, S1E), but accumulates along the polarity of PCP in the *Xenopus* neural plate. Accumulated Wnt11 is co-localized with core PCP components, suggesting local, reciprocal interaction. Interestingly, in the presence of core PCP components, Wnt11 can induce its own polarization and that of core PCP components, even if Wnt11 is not provided as a gradient. (Fig. 3A and 8I). Thus, non-directional Wnt11 and core PCP components can autonomously generate cell polarity without any directional information and Wnt11 can be recognized as a soluble component that interacts tightly with a core PCP component, i.e., a “soluble” core PCP component. Based on these results, we propose a model in which PCP is self-organized in a bottom-up manner through local and reciprocal interaction between Wnt11 and core PCP components, by forming asymmetric complexes (Fig. 8I).

We further discovered molecular mechanisms by which PCP emerges through local and reciprocal interactions between Wnt11 and core PCP components. We showed that Wnt11, which can induce Vangl2 phosphorylation, accumulates pVangl2 on the same cell membranes, and conversely Vangl2 phosphorylation promotes co-accumulation of pVangl2 and Fzd7 with Wnt11 (Fig. 6D-G). Thus, phosphorylated Vangl2 and Wnt11 accumulate one another in a mutual regulatory loop, accounting for the polarized localization of Wnt11 (Fig. 1D and 1E) and its requirement for Vangl2 (Fig. 1F and 1G). We propose that continuation of this regulatory loop increases local concentrations of Wnt11, Fzd7, and Vangl2, biasing the concentrations of these molecules on the cell membrane. In addition to this loop, we showed that two other mutual regulatory loops, in which each component reciprocally activates the other, are involved. One is a cis-interaction loop between non-phosphorylated Vangl2 and Pk3, which does not necessary involve Wnt11 (Fig. 7). The other is a trans-interaction between phosphorylated and non-phosphorylated Vangl2, located on opposing cell membranes (Fig. 8). Since these two regulatory loops and the amplification loop of Wnt11-Fzd7-pVangl2 are closely correlated, the two cis-interactions are further enhanced when they face each other to form a trans-interaction (Fig. 8). Thus, we propose that cell polarity can emerge by an amplification mechanism based on combinatorial interactions of these three positive regulatory loops.

The importance of cell-bridging interactions of core PCP components, including Vangl/Stbm, Fzd/Fz and Celsr/Fmi, for PCP has been reported in both mice and *Drosophila* (*32–34, 45*). Furthermore, our study demonstrates that the intercellular interaction between phosphorylated and non-phosphorylated Vangl2 is important in terms of modulating polarity of adjacent cells. Since phosphorylation of Vangl in vertebrates (and Vang/Stbm in *Drosophila*) has been described as an important event in PCP (*27, 40–42*), it is an interesting question whether the intercellular interaction mediated by phosphorylated and non-phosphorylated Vangl2 is also involved in many aspects of PCP formation, especially in *Drosophila* wing disc, where Wnt is not essential for PCP formation.

Since Wnt proteins co-assemble on HSPG clusters on the cell membrane (*35*), interactions between Wnt and HSPGs may influence accumulation of core PCP components and Wnt11 observed in this study. Interestingly, when Wnt ligands activate Wnt/β-catenin signaling, it induces assembly of signalosome components, including Fzd, on the cell membrane. Co-accumulations of Wnt ligands and Frizzled are also observed in Wnt-dependent cell polarization of *C. elegans* (*46*). Similarly, Dishevelled (Dvl), another signalosome component in Wnt/β-catenin signaling, is also accumulated by Wnt in the context of PCP (*47, 48*). Thus, we speculate that some machinery involved in signalosome formation may be used in Wnt-dependent assembly in PCP formation.

## Acknowledgements

We thank Drs. David Strutt and Helen Strutt for discussion; Dr. Sergei Y. Sokol for discussion, cDNAs of GFP-Pk3 and HA-Vangl2; Dr. Kazuhiro Aoki for cDNAs of mTagBFP2, KRasCT; Dr. Yusuke Miyanari for mRuby2 cDNA: Dr. Shinichi Higashijima for technical support, and Dr. Steven D. Aird for editing the manuscript. This work was supported in part by following programs: KAKENHI from the Japan Society for Promotion of Science (18H02454, 21H02498, 24111002, 17H05782, and 19H04797 to ST and 15K14532, 18K14720, 22H02637/23K23900, and 23H04930 to YM), PRESTO from Japan Science and

Technology Agency (JPMJPR194B to YM). Additional support also came from grants from National Institutes of Natural Sciences (NINS Joint Research Program to ST and YM).

## Data and materials availability

All codes and materials are available to academic researchers upon request.

