## Supplementary Materials for "Emergence of cell polarity by reciprocal interactions between Wnt and core PCP components"

### Supplementary Materials (Mii et al.)

#### Materials and methods

##### *Xenopus* embryo manipulation and microinjection.

All experiments using *Xenopus laevis* were approved by the Institutional Animal Care and Use Committee, National Institutes of Natural Sciences or the Animal Experimentation Committee, Kyoto University. Handling of *X. laevis* and microinjections were performed using standard methods (*49*). In brief, female *Xenopus* were injected with gonadotropin (ASKA Pharmaceutical) to obtain unfertilized eggs, which were then fertilized with testis homogenates. Fertilized eggs were de-jellied in 4% cysteine (pH7.8) and incubated in 1/10x Steinberg's solution at 14-20 °C. Embryos were staged according to Nieuwkoop and Faber (*50*). All presented data are a representative of at least two independent experiments. Microinjections were performed on early-stage embryos (4-32 cell) in 3% Ficoll (Sigma-Aldrich))/modified Birth's solution (MBS). mRNAs used in microinjections were synthesized from pCSf107mT (*51*)-based plasmid DNAs using an mMessage mMachine SP6 kit (Invitrogen) and purified with an RNeasy Micro kit (QIAGEN). Morpholino oligos (MOs) were used in knockdown experiments. Unless specified in Figure Legends, embryos were observed at Stage (St.) 14. Amounts of injected mRNAs/MOs are described in Figure Legends.

##### Morpholino oligos (MOs).

The following MOs were used in this study. *wnt11b,* CCAGTGACGGGTCGGAGCCATTGGT (*52*); *vangl2*, GAGTACCGGCTTTTGTGGCGATCCA (*53*) and CGTTGGCGGATTTGGGTCCCCCCGA (*54*) *fzd7*, CCAACAAGTGATCTCTGGACAGCAG (*55*) and GCGGAGTGAGCAGAAATCGGCTGAT (*56*). Standard (std) MO (CCTCTTACCTCAGTTACAATTTATA) was used as a negative control.

##### Fluorescent image acquisition.

Fluorescent images were acquired with a spinning disc confocal microscope (Andor Dragonfly 200 combined with an Olympus IX83, objective: UPLXAPO20X x20/NA0.8) or laser-scanning confocal microscope (Leica TSC SP8 system, objective: HC PL APO2 x40/NA1.10 W CORR water immersion).

##### Super-resolution microscopy.

Stimulated emission depletion (STED) microscopy was carried out with a Leica TCS SP8 STED 3X FALCON system (objective: HC PL APO 100x/1.40 OIL CS2) as previously described (*35*). Because wild-type embryos were destroyed by STED observation due to heat generated in pigment granules, albino embryos were used for analysis. These were fixed at St. 14 and stained with Alexa Fluor 488- and/or Alexa Fluor 568-labelled secondary antibody. For STED, a 592-nm laser was used for GFP and Alexa Fluor 488 and a 775-nm laser was used for Alexa Fluor 568. STED images were processed with "tau STED" function (LAS-X software, Leica) utilizing fluorescence lifetime analysis to enhance separation and localization of fluorescent peaks (*57*). Peak-to-peak distance was semi-automatically quantified using Fiji/ImageJ with the "FWHM line" plug-in (Dr. Lim Soon Yew John, in the ImageJ Mailing List).

#### Production of an anti-Wnt11 monoclonal antibody

We produced a mouse monoclonal antibody against *Xenopus* Wnt11 (clone #56-1), as described previously (*58*). Briefly, Recombinant full-length *Xenopus* Wnt11 was prepared as previously described (*35, 59*) and the antigen emulsion was injected into BALB/c mice. Treated mice were euthanized 21 days after injection, and lymphocytes were fused with SP2/0-Ag14 myeloma cells. After the cell fusion, culture supernatants were screened to confirm positive clones with a solid-phase enzyme-linked immunosorbent assay (ELISA). ELISA-positive clones were further screened by immunostaining of Wnt11-Myc overexpressing embryos.

#### Immunostaining.

*Xenopus* embryos were fixed with MEMFA (0.1M MOPS, pH 7.4, 2 mM EGTA, 1mM $MgSO_4$, 3.7% formaldehyde). For endogenous Wnt11 staining, fixed embryos were dehydrated with EtOH and rehydrated with TBT (1x TBS, 0.2% BSA, 0.1% Triton X-100). After replacement with TBT, fixed embryos were blocked with TBTS (TBT with 10% heat-immobilized FBS [70 ºC, 40 min]) for 1 h. For staining total Vangl2, antigen retrieval was performed at 95 ºC for 20 min in citrate-based Antigen Unmasking Solution (H-3300-250, Vector Laboratory) before blocking. Primary antibodies and their dilutions were as follows: anti-Wnt11 (mouse monoclonal IgG2b, in-house preparation, 1/10), anti-Vangl2 (HPA027043, Sigma, rabbit polyclonal IgG, 1/200), anti-Fzd7 (ab64636, Abcam, rabbit polyclonal IgG, 1/200), anti- Phospho-Vangl2-T78/S79/S82 (AP1206, ABclonal, rabbit monoclonal IgG, 1/1000), anti-C-Cadherin (6B6, Developmental Studies Hybridoma Bank, mouse monoclonal IgG1, 1/50), anti-β-catenin (C2206, Sigma, rabbit polyclonal IgG, 1/2000). Primary antibodies were diluted with Can Get Signal immunostain Immunoreaction Enhancer Solution A (NKB-501, TOYOBO) or B (NKB-601, TOYOBO) or TBTS. Secondary antibodies and their dilutions were as follows: goat polyclonal anti-mouse IgG Alexa Fluor 488 (A11001, Molecular Probes, 1/500), goat polyclonal anti-mouse IgG Alexa Fluor 647 (A21235, Molecular Probes, 1/500), goat polyclonal anti-rabbit IgG Alexa Fluor 488 (A11008, Molecular Probes, 1/500), goat polyclonal anti-rabbit IgG Alexa Fluor 555 (A21429, Molecular Probes, 1/500), goat polyclonal anti-rat IgG Alexa Fluor 555 (A21434, Molecular Probes, 1/500). See also Table S1 for antibodies. Secondary antibodies were diluted with TBTS. Embryos were incubated in antibody solution overnight at 4 ºC and washed with TBT 6 x 30 min after incubation with primary or secondary antibodies. Wheat germ agglutinin (WGA)-conjugated with Alexa Fluor 647 (W32466, Molecular Probes) was mixed in secondary antibody solution at dilution of 1/2000.

#### Image analysis.

All measurements of signal intensity were performed with Fiji/ImageJ (v1.54f). To obtain spatial intensity profiles of endogenous Wnt11 in the neural plate (Fig. 1C and S1E), intensity profiles were obtained from posterior to anterior along a 100-pixel width (= 151.67 μm) "Straight line". To evaluate effects of knockdown experiments on endogenous factors in the neural plate (Fig. 1F, G and S5D, E) and because basal signal intensity varied among embryos, normalized intensities of regions of interest (ROIs) were calculated by dividing intensities of ROIs by the averaged intensity of the uninjected side for each embryo. For Fig. S1B, signal intensity was normalized to the averaged Wnt11 intensity in std MO-injected embryos. Signal intensities of cell membranes were obtained by generating an intersection image of a cell segmentation mask (5 to 10-pixel width) and an image to be quantified using "Subtract" operation of the "Calculator Plus" plugin in Fiji/ImageJ. Segmentation masks were initially generated with "Cellpose" (using the model of "Cyto2" or "Cyto3") (*60*) based on membrane marker images and subsequently corrected by "TissueAnalyzer" (*61*) in the Fiji/ImageJ plugin. For measurement of signal intensities and correlation of factors in Fig. 5 to 6, 7, 8, S4, S5, S7, S8, and S9, ROIs of each cell boundary were obtained by manually tracing cell boundaries with lines 5- or 10-pixels using an iPad (Apple). Intensity profiles were collected with in-house Fiji/ImageJ macros, and averaged intensities and correlation coefficients among factors were calculated with in-house Python codes.

For quantification of polarity angles and polarity magnitude in Fig. 1, 2, 3, 8, and S3, cell segmentation images were generated as described above. Polarity angles and magnitudes were quantified by "QuantifyPolarity2.0" with the PCA method (*43*). For polarity angles of GFP-Pk3 (Fig. 2E) and mRuby2-Pk3 (Fig. 4N), embryos with clearly separable Wnt- and Pk3-expressing regions and with apparent membrane localization of Pk3 were selected (*24*).

To evaluate endosomes (Fig. 6H), intracellular vesicles were identified from binarized Wnt11-BFP images using the "Analyze Particles" function in Fiji/ImageJ (see also Fig. S9E). Image binarization was performed using the "Triangle" method. Vesicle size was thresholded at 0.04 $\mu m^2$.

To calculate concentration indices for evaluating signal accumulations (Fig. 4 and 6), the number of pixels whose intensity was higher than the mean intensity of the cell boundary was divided by the total number of pixels.

#### Statistical analyses.

Sample sizes were determined empirically. Unless specified in Figure Legends, statistical analyses were performed with the Wilcoxon rank-sum test adjusted for multiple comparisons with Holm's method.

#### Supplementary Figures

Fig. S1. Gene expression pattern of *wnt11* and specificity of anti-Wnt11 immunostaining.

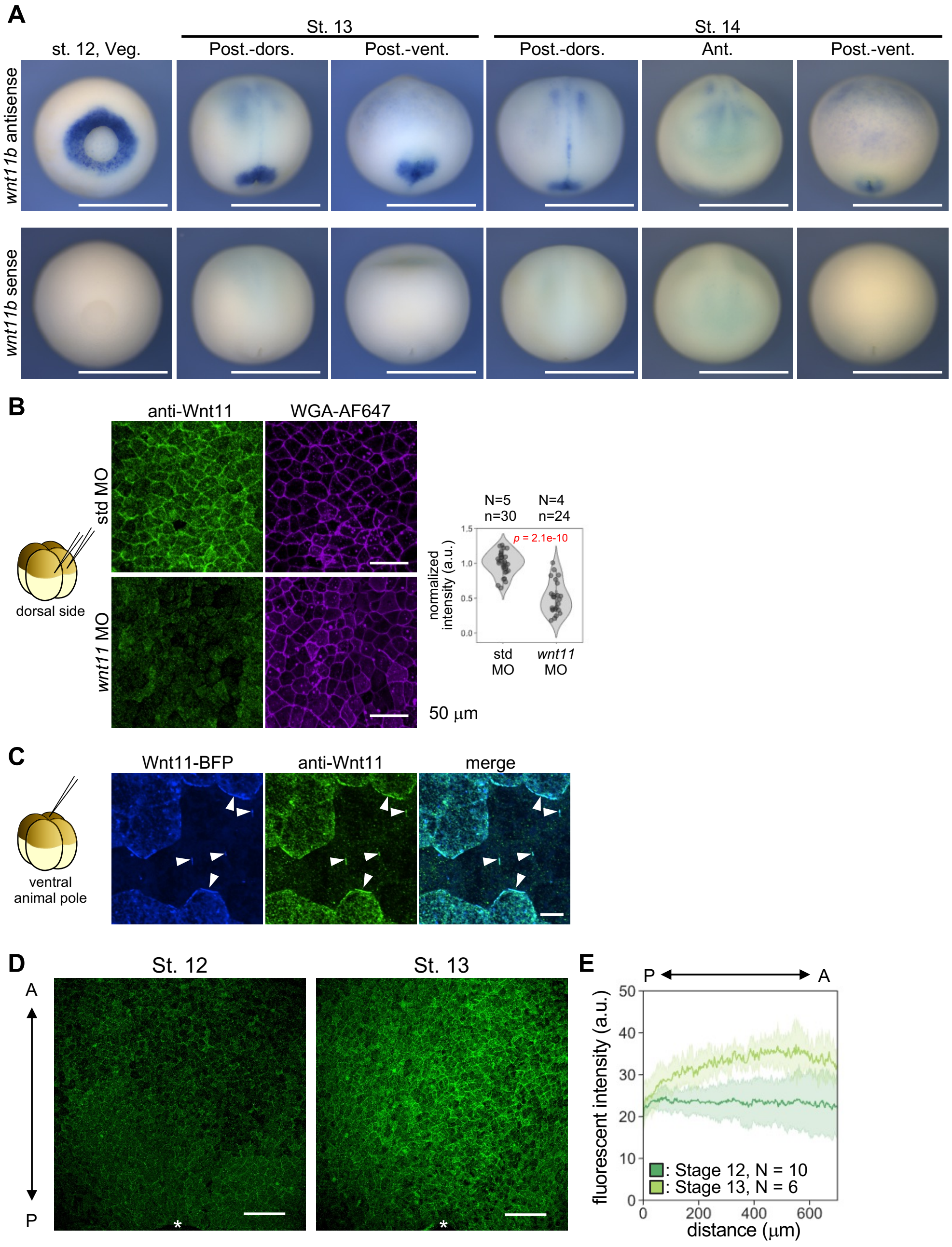


**A, Gene expression patterns of *wnt11b* visualized by whole-mount in situ hybridization.** Throughout Stages 12-14, *wnt11b* mRNA is highly localized at the posterior region, near the blastopore. Weaker expression is also observed in the anterior neural plate at Stages 13 and 14.

**B, Reduction of anti-Wnt11 staining by knockdown of *wnt11*.** Alexa Fluor 647-conjugated wheat germ agglutinin (WGA-AF647) was used as a membrane marker. Numbers of embryos (N) and numbers of subregions (n, squares of 98 µm sides) are as indicated. MOs were injected as illustrated. Amounts of MOs (ng/embryo): *wnt11* MO, 42.7; std MO, 41.6 (both are 5 pmol).

**C, anti-Wnt11 staining with overexpression of mTagBFP2-tagged Wnt11 (Wnt11-BFP, W11B).** Anti-Wnt11 staining overlapped with localization of Wnt11-BFP, including accumulation and puncta at cell boundaries (arrowheads). *wnt11-BFP* mRNA (1.0 ng/embryo) was injected as illustrated.

**D, Immunostaining of endogenous Wnt11 in *Xenopus* embryos at earlier stages.** Posterior-dorsal view. The anterior-posterior (AP) axis is as indicated. Asterisks indicate positions of the dorsal edge of blastopore.

**E, Spatial profile of Wnt11 in the dorsal/neural plate region at earlier stages.** Wnt11 did not form a concentration gradient from posterior to anterior in dorsal ectoderm of St. 12 or in the neural plate of St. 13. Fluorescent intensities were obtained from line plots 152 µm wide at the center of the dorsal/neural plate region. Error ranges are given as standard deviations (s.d.). Numbers of embryos (N) are as indicated.

Scale bars: 1 mm (A), 50 µm (B), 10 µm (C), 200 µm (D).

Fig. S2. Activity of BFP-tagged and membrane-tethered Wnt11 for PCP formation.

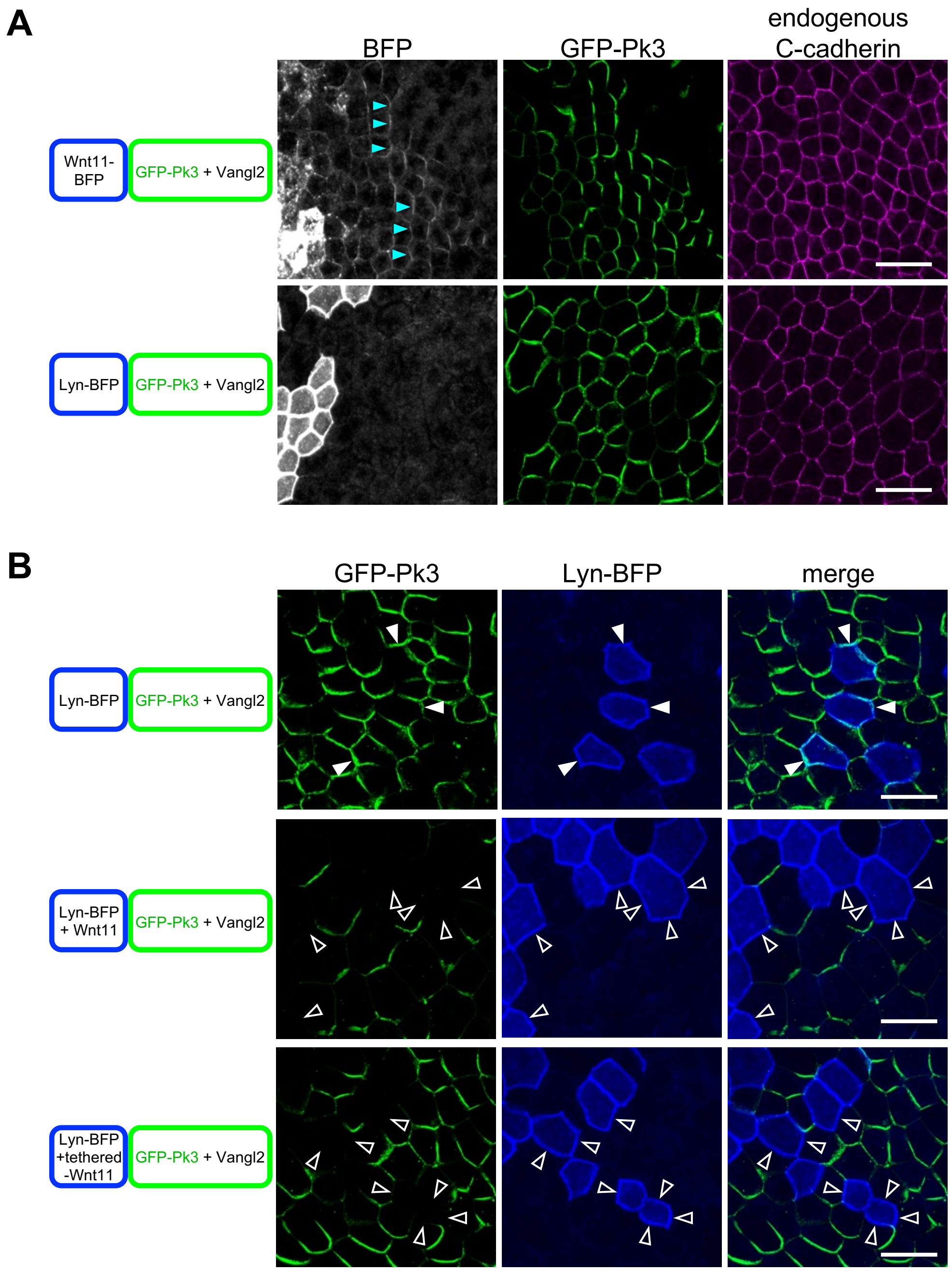


**A, Reconstructed PCP (rPCP) formation with Wnt11-BFP in the animal cap region.** Wnt11-BFP can polarize GFP-Pk3 co-expressed with HA-Vangl2 (upper panels), whereas a membrane tracer, Lyn-BFP cannot (bottom panels). As with endogenous Wnt11 in the neural plate, Wnt11-mTagBFP2 exhibits polarized localization, co-localized with GFP-Pk3 (asterisks). A membrane marker, C-cadherin is uniformly distributed regardless of polarization of GFP-Pk3. **B, Cell membrane localization of GFP-Pk3 was reduced by both wild-type and membrane-tethered Wnt11 expressed in adjacent cells.** Membrane localization of GFP-Pk3 was reduced by both wild-type (WT) and membrane-tethered-Wnt11 expressed in adjacent cells (open arrowheads), but not by Lyn-BFP (closed arrowheads). Note that reduction of GFP-Pk3 occurred at cell boundaries in various directions, suggesting contact-dependent effects of Wnt11-source cells. mRNAs were injected into ventral animal blastomeres at the 32-cell stage as illustrated. Amounts of mRNAs (pg/embryo): *wnt11-BFP*, 500; *lyn-BFP*, 50; *GFP-pk3*, 200; *HA-vangl2*, 100; *wnt11*, 500; *tethered-wnt11*, 500. Scale bars, 50 μm.

Fig. S3. Exogenous Vangl2 stabilizes Fzd7 protein on membranes.

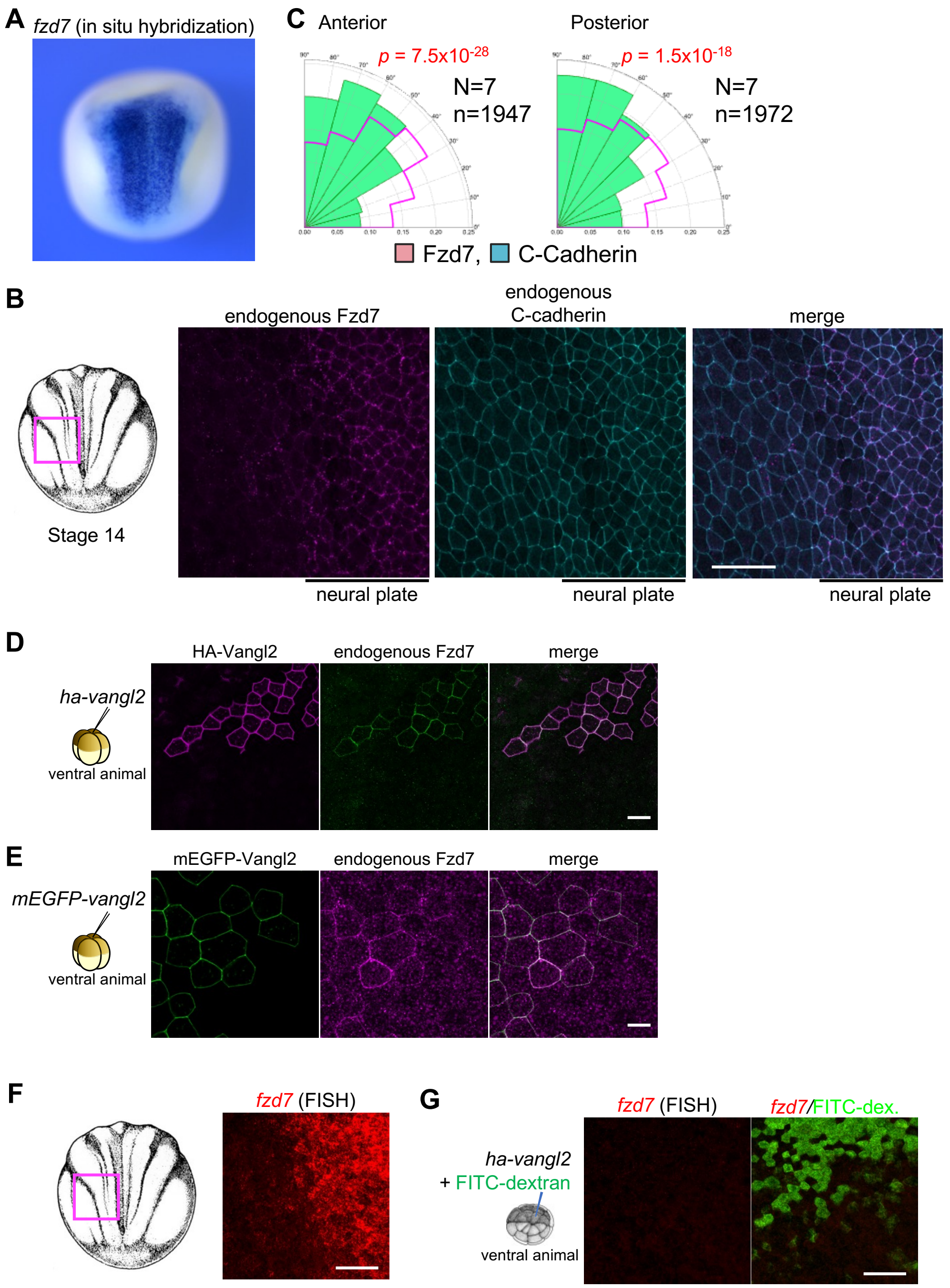


**A, Gene expression patterns of *fzd7* mRNA visualized by whole-mount in situ hybridization.** *fzd7* is highly expressed in the neural plate. **B, Membrane localization of endogenous Fzd7 was specifically observed in the neural plate.** A boundary region of the neural plate was observed as illustrated. C-cadherin was stained as a membrane marker. **C, Quantification of the polarity angle of endogenous Fzd7 localization in the neural plate.** Anterior and posterior halves of neural plates were analyzed. In both regions, Fzd7 was significantly polarized toward 90º (corresponding to localization on the medio-lateral boundaries) compared to C-cadherin. **D, E, Overexpression of Vangl2 in the animal cap region increased membrane localization of Fzd7.** HA-Vangl2 (D) or mEGFP-Vangl2 (E) was ectopically expressed. Both increased membrane localization of endogenous Fzd7. **F, G, Fluorescent *in situ* hybridization (FISH) of *fzd7*.** (F) Boundary of the neural plate. Similar to normal in situ hybridization (A), *fzd7* transcripts were intensively detected in the neural plate. (G) Overexpression of HA-Vangl2 did not increase *fzd7* transcripts in the animal cap region, suggesting that the increase of Fzd7 by Vangl2 overexpression is probably due to stabilization of Fzd7 protein, but not due to an increase of *fzd7* transcripts. Samples were stained and visualized in a side-by-side manner to equalize sensitivity for *fzd7* transcripts in F and G.

mRNAs were injected as illustrated. Amounts of mRNAs (pg/embryo): *HA-vangl2*, 50 (D), 100 (G); *mEGFP-vangl2*, 100. Scale bars, 100 µm (B, F, G), 40 µm (D), 20 µm (E).

Fig. S4. Dose-dependent reduction and biphasic polarization of GFP-Pk3 by co-expressed Wnt11.

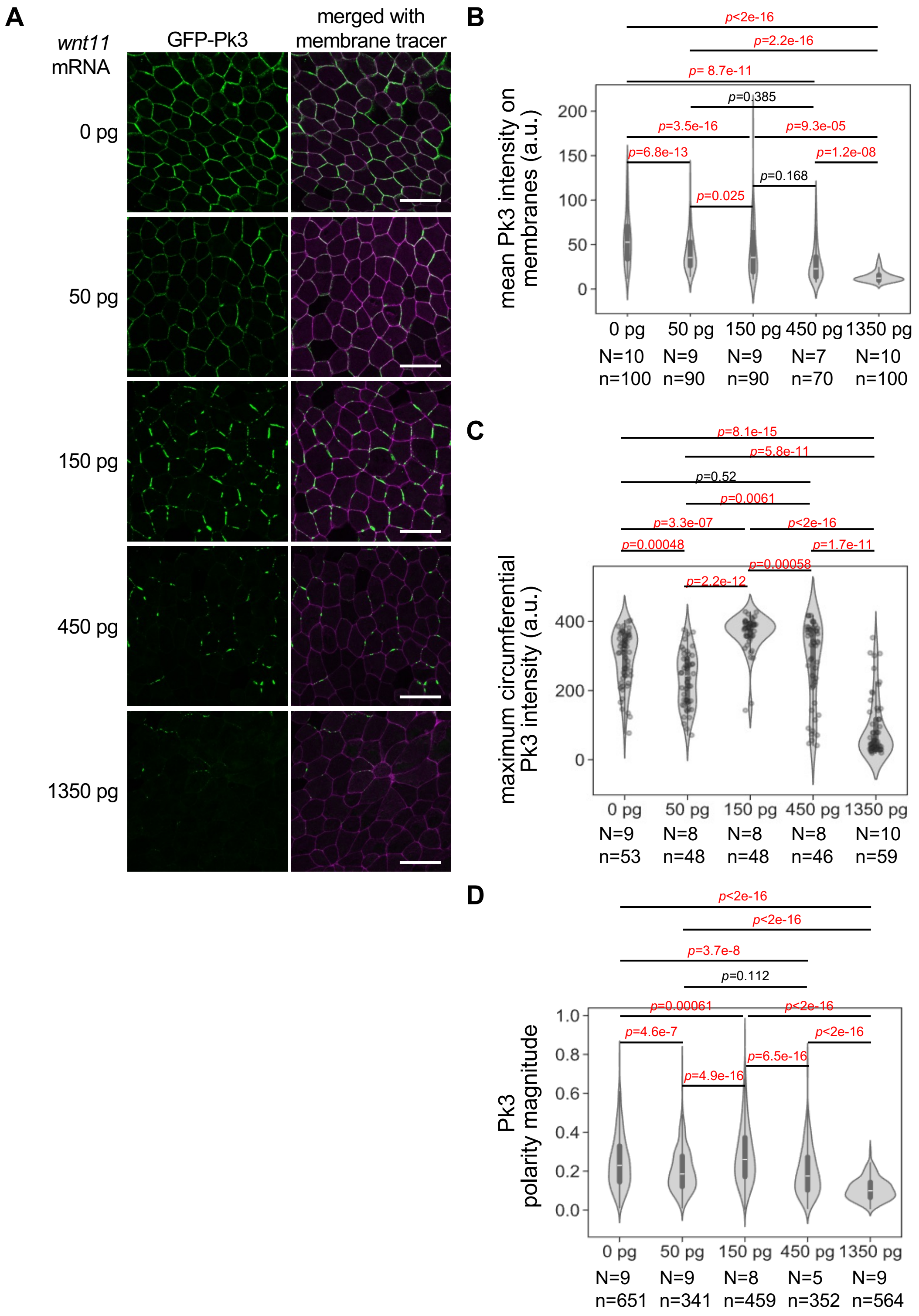


**A, Localization of GFP-Pk3. B, Quantification of mean intensity of GFP-Pk3 on cell membranes.** GFP-Pk3. Numbers of embryos (N) and numbers of cell boundaries (n) are as indicated. Wnt11 reduced GFP-Pk3 on membranes in a dose-dependent manner. **C, Circumferential maximum intensities of GFP-Pk3.** Unlike the mean intensity, maximum intensity was highest with an intermediate dose of wnt11 mRNA (150 pg/embryo). Numbers of embryos (N) and number of cells (n) are as indicated. **D, Quantification of polarity magnitude of GFP-Pk3.** As with the maximum intensity, polarity magnitude was also highest with an intermediate dose of wnt11 mRNA (150 pg/embryo). Numbers of embryos (N) and number of cells (n) are as indicated.

mRNAs were injected into a ventral animal blastomere at the 16-cell stage. mRNAs (pg/embryo): *wnt11*, as indicated; *GFP-pk3*, 200; *HA-vangl2*, 100; *mRuby2-kras*, 67. Scale bars, 50 µm.

281 Fig. S5. Specificity of anti-phospho Vangl2 (pVangl2) staining.

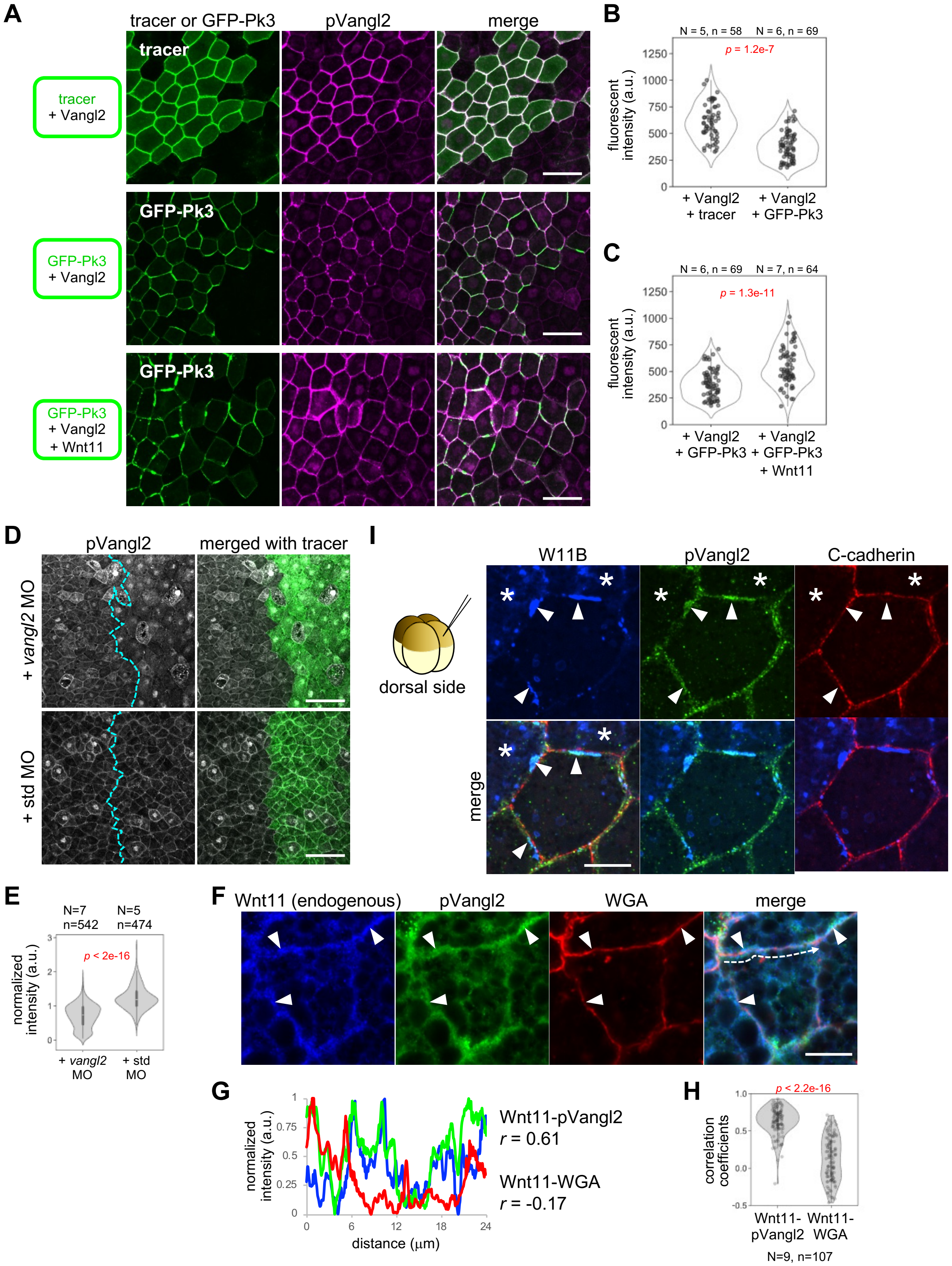

**A-C, pVangl2 staining in overexpression of Vangl2, GFP-Pk3, and Wnt11.** (A) mRNAs were injected into the animal pole region of a ventral blastomere at the 4-cell stage. Overexpression of Vangl2 in the animal cap region increased pVangl2 staining, compared to tracer-negative cells. mEGFP-KRasCT was used as a tracer. Co-expression of GFP-Pk3 reduced pVangl2 staining, compared to co-expression of the tracer, consistent with biochemical analyses with Xenopus embryos (44). Further addition of Wnt11 to co-expression of Vangl2 and GFP-Pk3 increased pVangl2 staining, suggesting that Wnt11 can induce phosphorylation of Vangl2. (B, C) Quantification of staining intensity of pVangl2. A single data point corresponds to an average intensity of a line plot on a cell boundary between tracer/GFP-Pk3-positive and -negative cells. Numbers of embryos (N) and numbers of cell boundaries (n) are as indicated. For (+ Vangl2, + GFP-Pk3), the same data are presented in B and C.

**D, E, pVangl2 staining is reduced by vangl2 MO in the neural plate.**
MOs (14 ng/embryo) were injected into a dorsal blastomere at the 4-cell stage, targeting the future neural plate. FITC-dextran was used as a tracer. Intensity of pVangl2 on cell membranes was quantified in injected and uninjected sides, and normalized by the mean value of the uninjected side in each embryo.

**F, G, Co-localization of endogenous Wnt11 and pVangl2 at subcellular resolution in the neural plate.** Wnt11 and pVangl2 are frequently co-localized in a punctum-by-punctum manner (arrowheads) but not with WGA (membrane marker).

E, Intensity profiles of Wnt11, pVangl2 and WGA at a cell boundary. Normalized intensities along the dashed arrow (D) are plotted.

**H, Quantification of correlation coefficients.** Correlation coefficients between Wnt11 and pVangl2 are significantly higher than those between Wnt11 and WGA. 107 cell boundaries from 9 embryos were analyzed. Statistical analysis was performed with the Wilcoxon signed rank test with continuity correction.

**I, Overexpression of Wnt11-BFP (W11B) caused co-accumulation of W11B and pVangl2 in the neural plate.**
Overexpression of W11B in the neural plate often caused self-accumulation at cell boundaries (arrowheads). At these accumulations, W11B co-localized with pVangl2, but not C-cadherin. W11B-expressing cells are indicated with asterisks.

Amounts of mRNAs: *vangl2*, 100; *gfp-pk3*, 200; *mEGFP-KRasCT*, 50; *wnt11*, 250; *wnt11-BFP* 1.0 ng/embryo.

Scale bars, 50 μm (A),100 μm (D) 10 μm (I). Numbers of embryos (N) and numbers of cells (n) are as indicated.

Fig. S6. Resolution of STED microscopy analyzed by peak-to-peak distance of membrane-tracers.

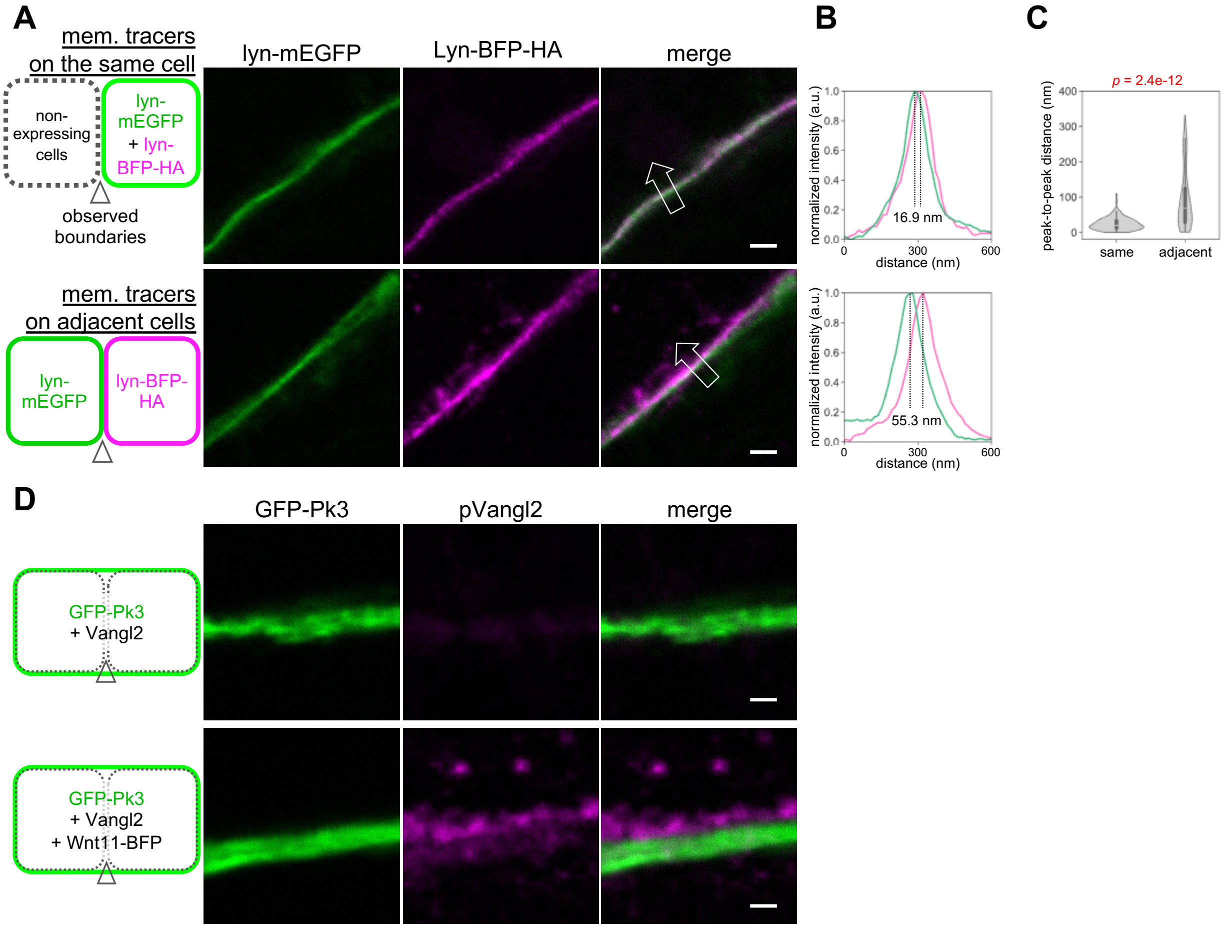


**A-C, STED microscopy can resolve different membrane tracers expressed on adjacent cells.** (A) Lyn-mEGFP and Lyn-BFP-3HA were expressed on the same cells or on adjacent cells in the animal cap region of *Xenopus* embryos. (B, C) Examples of quantification of peak-to-peak distance between the two membrane tracers. Line plots at positions indicated by arrows in A are shown. For a single line, a width of 20 px = 284 nm was measured. (C) Peak-to-peak distance (PPD). A significant difference was observed, indicating that membrane proteins on adjacent cells can be resolved by our STED observations. Data from 4 embryos, 23 cell boundaries, 119 lines (same cell) and 4 embryos, 21 cell boundaries, 115 lines (adjacent cells) are presented.

**D, pVangl2 staining observed with STED microscopy.** In the absence of overexpressed Wnt11-BFP, staining of pVangl2 was weak. In the presence of overexpressed Wnt11-BFP, pVangl2 staining was strong and clearly separated from GFP-Pk3.

mRNAs were injected into animal pole regions of ventral blastomeres at the 4-cell stage, as indicated. Amounts of mRNAs (ng/embryo): *lyn-mEGFP*, 0.10; *lyn-BFP-HA*, 0.10; *GFP-pk3*, 0.20; *vangl2*, 0.10; *wnt11-BFP*, 1.0. Scale bars, 500 nm (A, D).

Fig. S7. Vangl2 stabilizes Fzd7 on the same membrane in the absence of Wnt11.

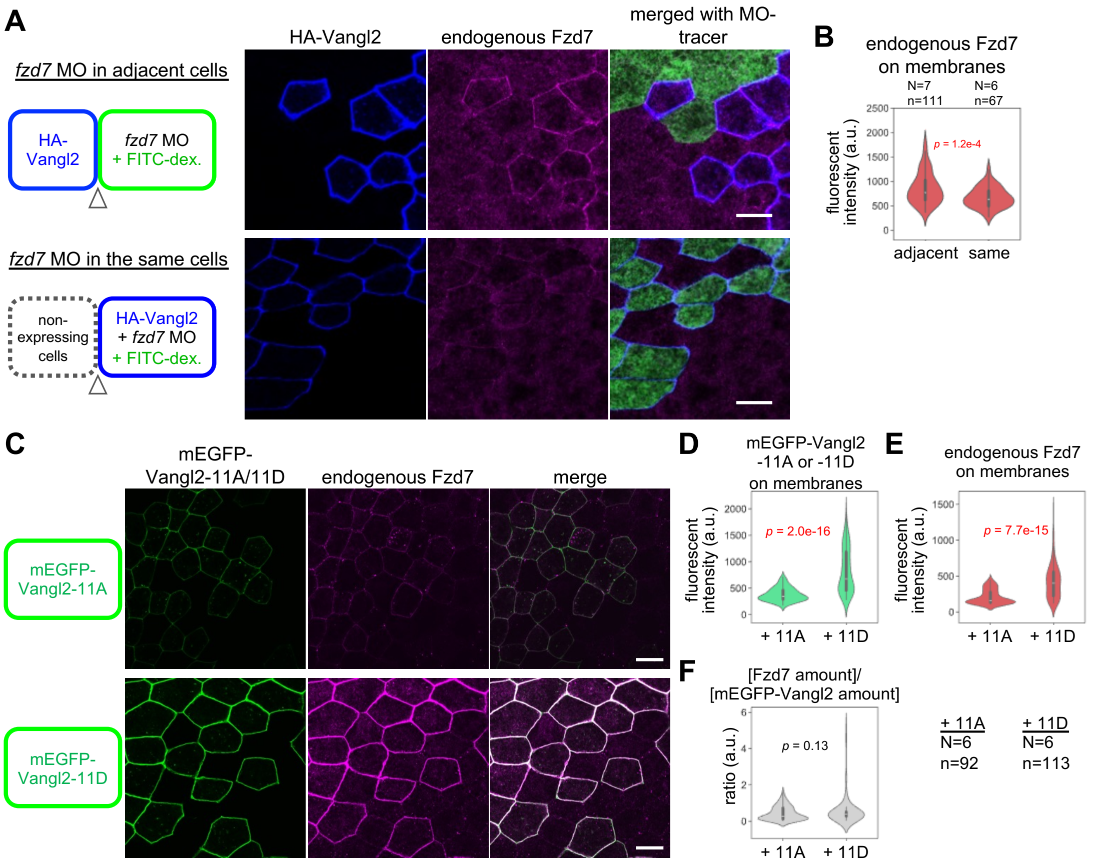


**A, B, Vangl2 increases membrane localization of Fzd7 on the same membrane.** *HA-vangl2* mRNA and *fzd7* MO were injected into ventral animal blastomeres at the 8-16-cell stage as illustrated. (A) Overexpression of HA-Vangl2 increased membrane localization of endogenous Fzd7 (Supplementary Fig. S2). *fzd7* MO in cells adjacent to HA-Vangl2-expressing cells did not affect Fzd7 on membranes (top panels), but *fzd7* MO in the same cells as HA-vangl2 reduced Fzd7 on membranes (bottom panels). (B) Quantification of fluorescent intensity of endogenous Fzd7 on membranes. To confirm which side of Fzd7 was knocked down, cell boundaries between HA-Vangl2-expressing cells and MO-tracer-containing cells, and cell boundaries between HA-Vangl2-expressing cells and non-expressing cells are quantified for "*fzd7* MO in adjacent cells" and "*fzd7* MO in the same cells", respectively. Numbers of embryos (N) and numbers of cell boundaries (n) are as indicated.

**C-F, Phospho-deficient and -mimetic mutants of Vangl2 can increase membrane localization of Fzd7 proportionally to the amount of Vangl2 protein on membranes.** mEGFP-Vangl2-11A or -11D was overexpressed in the animal cap region. As with wild-type Vangl2, phospho-deficient (11A) and -mimetic (11D) mutants increased endogenous Fzd7 (C). To equalize quantification of proteins on membranes, cell boundaries between mEGFP-Vangl2-11A/D-expressing cells and non-expressing cells are quantified. Amounts of Vangl2 itself (D) and endogenous Fzd7 (E) were significantly higher with 11D-overexpression than those with 11A-overexpression. However, the ratio of [Fzd7 amount]/[mEGFP-Vangl2 amount] did not differ significantly between 11A and 11D (F), suggesting that the increase of Fzd7 is proportional to the amount of Vangl2 protein on membranes, regardless of phosphorylation states. Data of 92 cell boundaries from 6 embryos (+ 11A) and 113 cell boundaries from 6 embryos (+ 11D) are presented in D-F.

Amounts of mRNAs/MOs (ng/embryo): *HA-vangl2*, 0.050; *mEGFP-vangl2* (*-11A*, *-11D*); 0.10; *fzd7* MOs, 3.5. Scale bars, 20 μm.

Fig. S8. Vangl2 is required for membrane localization of Fzd7 on the same membranes in the absence of Wnt11.

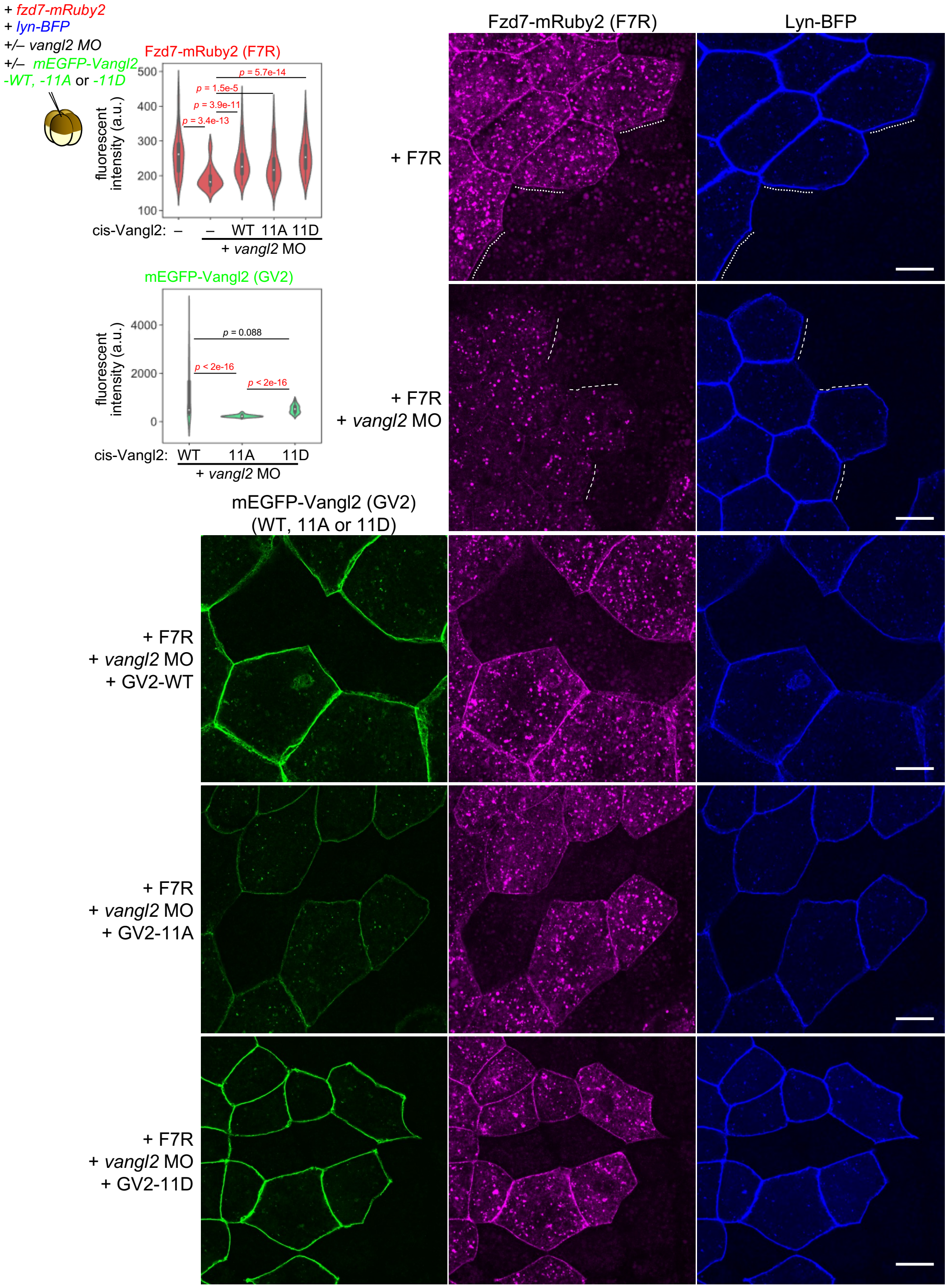


mRNAs and *vangl2* MO were injected into the animal pole of a ventral blastomere as illustrated. (Pictures) Fzd7-mRuby2 (F7R) exhibited membrane localization (dotted lines). Co-injected vangl2 MO reduced membrane localization of F7R (dashed lines). Additional expression of either the wild-type (WT) mEGFP-Vangl2 (GV2), -11A or -11D can rescue membrane localization of F7R on the same membrane (see cell boundaries between tracer-expressing and non-expressing cells). (Graphs) Cell boundaries between tracer-expressing and non-expressing cells were quantified for amounts of F7R and GV2 on the same membranes. Quantification indicates that GV2-WT and GV2-11D fully rescue membrane localization of F7R, whereas GV2-11A does so partially. In this experiment, translation of F7R is equalized because it completely depends on injected mRNA. Thus, Vangl2 is likely to stabilize Fzd7 on the same membrane, regardless of phosphorylation state. 75 cell boundaries from 5 embryos for each experimental group were quantified.

Amounts of mRNAs/MOs (ng/embryo): *fzd7-mRuby2*, 0.10; *mEGFP-vangl2* (WT, *-11A* and *-11D*), 0.10; *lyn-BFP*, 0.025; *vangl2* MO, 14. Scale bars, 20 µm.

Fig. S9. Effects of Wnt11 on Vangl2 and Fzd7 on the same membranes and endosomal removal of Pk3.

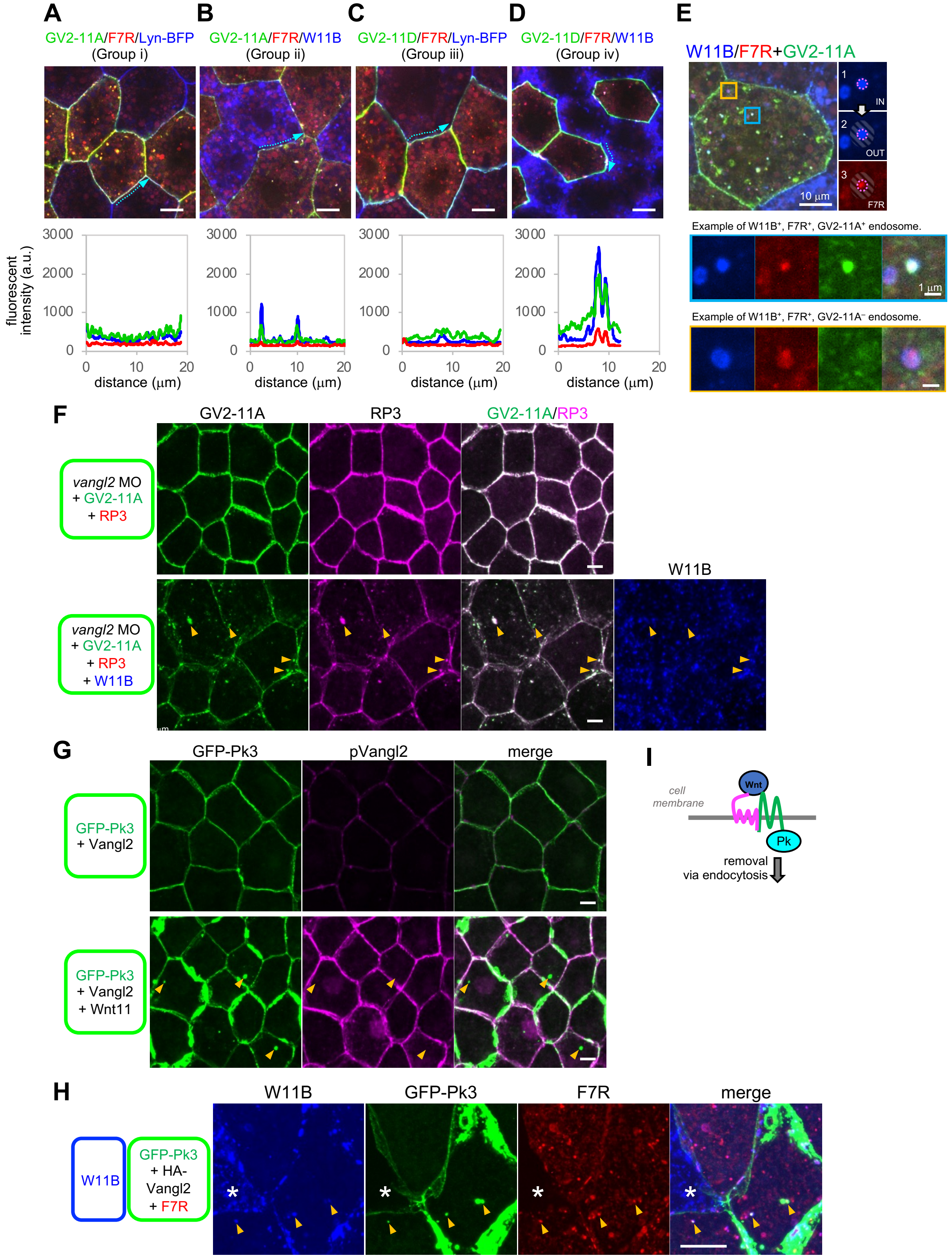

**A-D, Intensity profiles of each signal along outer cell boundaries shown by dotted arrows (blue, W11B; green, GV2; red, F7R)**. W11B, GV2 and F7R showed co-accumulation when W11B was co-expressed. Accumulation of these proteins is much higher with GV2-11D than with GV2-11A. Images are the same as Fig. 6D.

**E, Analysis of W11B-containing endosomes.** W11B-positive puncta inside receiving cells must be internalized from the extracellular space and were identified as endosomes by thresholding the image of W11B. Inside was defined as $ROI_{IN}$ (1). Then, surrounding area was defined as $ROI_{OUT}$ (2). For GV2-11A/D and F7R, presence or absence was determined with mean intensity of ROIs as following (3); $ROI_{IN} > ROI_{OUT}$: positive. $ROI_{IN} < ROI_{OUT}$: negative.

**F, Reduction of mRuby2-Pk3 (RP3) from cell membranes and endosomal internalization with GV2-11A by Wnt11.** Membrane localization of RP3 was decreased and endosome-like puncta (arrowheads) were increased by co-expression of W11B.

**G, Endosome-like puncta of GFP-Pk3 (arrowheads) did not overlap pVangl2 staining.**

**H, GFP-Pk3 is endocytosed with F7R and W11B.** Because W11B is expressed in adjacent cells, W11B-positive puncta inside GFP-Pk3-, HA-Vangl2- and F7R-expressing cells must be internalized by endocytosis.

**I, Schematic illustration of endosomal removal of Pk3-non-phosphorylated Vangl2-Fzd7 cis-complexes with Wnt11.**

mRNAs were injected into the animal pole region of ventral blastomeres at the 4-cell stage, as illustrated. Amounts of mRNAs/MOs (ng/embryo): *mEGFP-vangl2-11A*, 0.10 (A-E), 0.050 (F); *mEGFP-vangl2-11D*, 0.10; *fzd7-mRuby2*, 0.050 (A-D), 0.10 (E) 0.040 (H); *wnt11*, 0.25; *wnt11-BFP*, 0.50; *lyn-BFP*, 0.050; *GFP-pk3*, 0.20 (G), 0.10 (H); *mRuby2-pk3*, 0.10; *vangl2*, 0.10; *HA-vangl2*, 0.050; *vangl2* MO, 42 (A-D), 21 (F). Scale bars, 10 μm (A-D, E (top panel), F and G), 1 μm (E (middle and bottom panels)).

Fig. S10. Images of individual channels of trans-assembly of core PCP components by Wnt11.

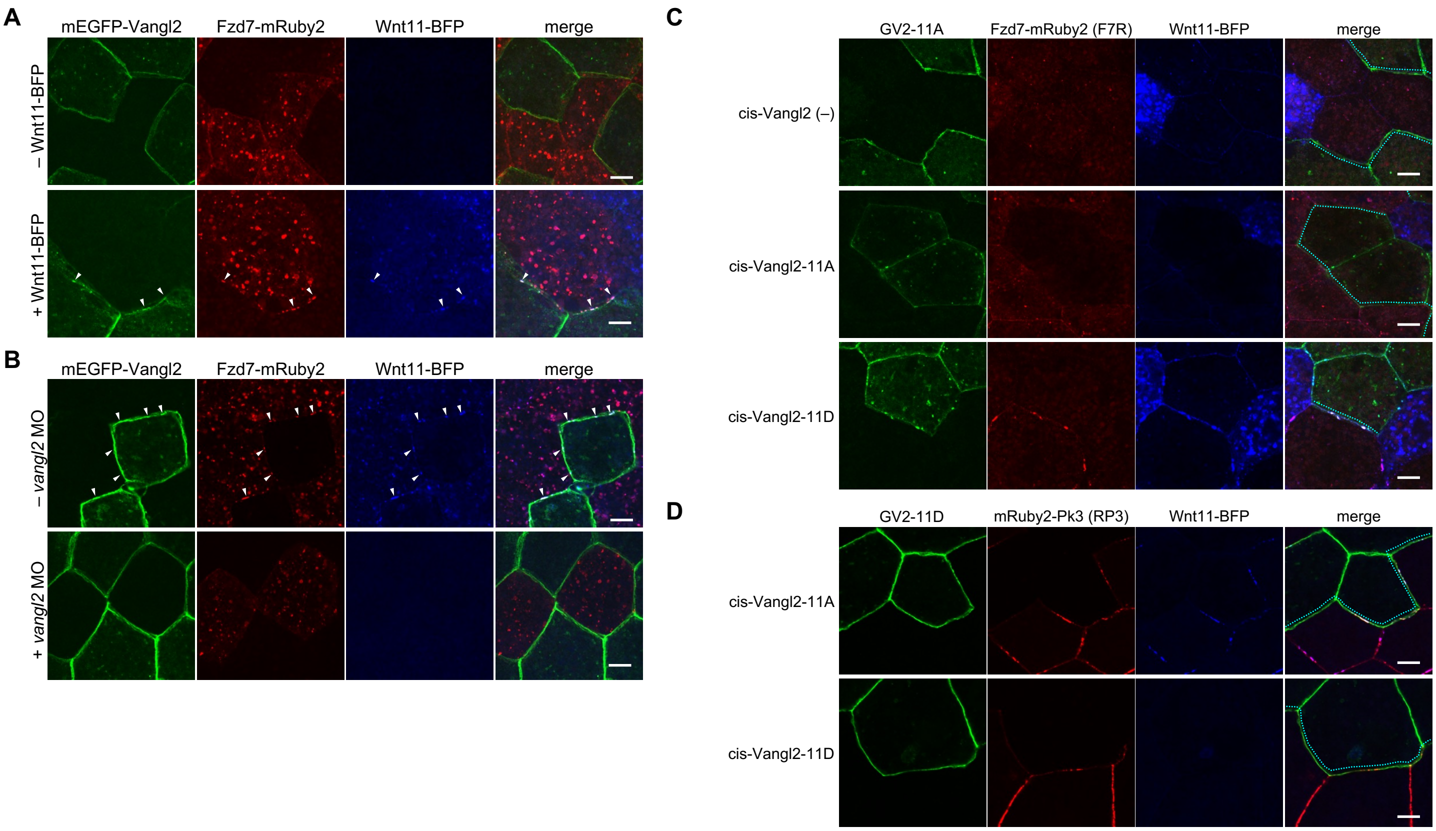


Images of individual channels corresponding to Fig. 8A (A), 8C (B), 8E (C) and 8G (D). Scale bars, 10 μm.

#### Table S1. Key resources.

| Reagent type or resource | Designation | Source or reference | Identifiers | Additional information | RRID |
|---|---|---|---|---|---|
| Antibody | Anti-HA (3F10 rat IgG1) | Roche | Cat no. 11867431001 | 1:2000 | AB_390919 |
| Antibody | Anti-XWnt11 (mouse monoclonal IgG2b) | this study | Clone: #56-1 | 1:10 | |
| Antibody | Anti-C-Cadherin (mouse monoclonal IgG1) | Developmental Studies Hybridoma Bank | Clone: 6B6 | 1:50 | AB_528113 |
| Antibody | Anti-b-catenin (rabbit polyclonal IgG) | Sigma | Cat no. C2206 | 1:2000 | AB_476831 |
| Antibody | Anti-Vangl2 (rabbit polyclonal IgG) | Sigma | Cat no. HPA027043 | 1:200 | AB_10601709 |
| Antibody | Anti-Phospho-Vangl2-T78/S79/S82 | ABclonal | Cat no. AP1206 | 1:200 | |
| Antibody | Anti-Fzd7 (rabbit polyclonal IgG) | Abcam | Cat no. ab64636 | 1:200 | AB_1640522 |
| Antibody | anti-mouse IgG Alexa Fluor 488 (goat) | Molecular Probes | Cat no. A11001 | 1:500 | AB_2534069 |
| Antibody | anti-mouse IgG Alexa Fluor 647 (goat) | Molecular Probes | Cat no. A21235 | 1:500 | AB_2535804 |
| Antibody | anti-rabbit IgG Alexa Fluor 488 (goat) | Molecular Probes | Cat no. A11008 | 1:500 | AB_143165 |
| Antibody | anti-rabbit IgG Alexa Fluor 555 (goat) | Molecular Probes | Cat no. A21429 | 1:500 | AB_2535850 |
| Antibody | anti-rat IgG Alexa Fluor 555 (goat) | Molecular Probes | Cat no. A21434 | 1:500 | AB_2535855 |
| Lectin | Wheat germ agglutinin (WGA)-conjugated with Alexa Fluor 647 | Molecular Probes | Cat no. W32466 | 1:500 | |
| Software, algorithm | QuantifyPolarity2.0 | (43) | | | |
| Software, algorithm | Cellpose | (60) | | | |
| Software, algorithm | TissueAnalyzer | (61) | | | |
| Software, algorithm | MATLAB | Mathworks | | | |
| Software, algorithm | ImageJ/Fiji (v1.54f) | | | | |
| Software, algorithm | RStudio | | | | |
| Software, algorithm | Spyder | | | | |

#### Movie S1. Time-lapse imaging of rPCP formation in the animal cap region.

Live-imaging of GFP-Pk3 and mRFP1 (tracer for Wnt11) during PCP formation (total duration: 9 hours, St. 10.25-14). One second in the movie corresponds to actual one hour.
